# Explaining the effects of distractor statistics in visual search

**DOI:** 10.1101/2020.01.03.893057

**Authors:** Joshua Calder-Travis, Wei Ji Ma

## Abstract

Visual search, the task of detecting or locating target items amongst distractor items in a visual scene, is an important function for animals and humans. Different theoretical accounts make differing predictions for the effects of distractor statistics. Here we use a task in which we parametrically vary distractor items, allowing for a simultaneously fine-grained and comprehensive study of distractor statistics. We found effects of target-distractor similarity, distractor variability, and an interaction between the two, although the effect of the interaction on performance differed from the one expected. To explain these findings, we constructed computational process models that make trial-by-trial predictions for behaviour based on the full set of stimuli in a trial. These models, including a Bayesian observer model, provided excellent accounts of both the qualitative and quantitative effects of distractor statistics, as well as of the effect of changing the statistics of the environment (in the form of distractors being drawn from a different distribution). We conclude with a broader discussion of the role of computational process models in the understanding of visual search.

## 1 Introduction

Animals and humans constantly engage in visual search, the process of detecting, locating or identifying target objects in an image or scene (Eckstein, 2011). The golden eagle looking for a hare, the hare looking for predators, and the human looking for bread in a supermarket are all examples of visual search. A great deal of research over more than 50 years has aimed to build a mechanistic understanding of visual search (Neisser, 1964; Treisman & Gelade, 1980; Estes & Taylor, 1964). Such an understanding would not just be important in its own right, but would contribute to our knowledge of the representations and algorithms used to perceive and act in the world. Additionally, it may have direct application in critical visual search situations, such as baggage scanning at the airport (Schwaninger, 2005).

Any satisfactory mechanistic account needs to explain key qualitative patterns in visual search data, such as those identified by Treisman and Gelade (1980). Treisman and Gelade (1980) suggested that there is a categorical difference between two kinds of search, which they called feature search and conjunction search. In feature search, the target can be distinguished from the distractor stimuli using a single feature such as colour. In conjunction search, there is no single feature which is present in the targets and absent in all distractors. Instead, two or more features are required to uniquely identify a stimulus as the target. Feature search was highly efficient: Increases in the number of stimuli in the display (the total number of targets and distractors) had little effect on the time taken to respond when a target was present. By contrast, efficiency in conjunction search was lower, and as set size increased, Treisman and Gelade (1980) found that response time increased markedly.

Duncan and Humphreys (1989) contested the idea of a dichotomy between feature and conjunction search and suggested that search efficiency varies along a continuum. They claimed that the similarity of the target to the distractors decreases search efficiency, variability of the distractors decreases search efficiency, and that these quantities interact such that the very hardest searches involved highly variable distractors which were very similar to the target. One of the key motivations for their departure from the idea of a strict dichotomy was that, over a series of experiments using a wide range of stimuli, they could not find a consistent set of properties which could be identified as features. Duncan and Humphreys’ (1989) account can still accommodate the findings of Treisman and Gelade (1980), if we claim that Treisman and Gelade (1980) only explored part of the stimulus space spanned by target-distractor similarity and distractor variability; the appearance of a dichotomy would then stem from the use of stimuli drawn from two distinct clusters in this space.

While Duncan and Humphreys (1989) made a valuable contribution in suggesting that performance likely lies on a continuum, their exploration of this claim was necessarily limited by the stimuli that they used. Using letters and joined lines, they could not parametrically vary properties of these stimuli along an easily quantified dimension. Therefore, it could be that Duncan and Humphreys (1989), while describing performance in a larger area of stimulus space than Treisman and Gelade (1980), missed areas of the space, along with distinctive qualitative effects.

Other visual search researchers have used stimuli that can easily be parametrically varied (Palmer, Ames & Lindsey, 1993; Palmer, 1994; Cameron, Tai, Eckstein & Carrasco, 2004; Ma, Navalpakkam, Beck, van den Berg & Pouget, 2011; Rosenholtz, 2001; Palmer, Verghese & Pavel, 2000). For example, Cameron et al. (2004) used oriented Gabor patches, with the target being defined as a Gabor of particular orientation. Such stimuli make it possible to operationalise target-distractor difference precisely as the difference between the target orientation and the mean distractor orientation, and to operationalise distractor variability as the variance of the distractors.

Parametric stimuli also allow us to apply formal models of behaviour and cognition (Ma et al., 2011; Rosenholtz, 2001; Palmer et al., 2000). Computational modelling could provide a simple unified explanation of the full range of patterns observed in behaviour, and could allow us to infer the precise mechanisms underlying visual search. Signal detection theory (SDT) is the leading framework for building such formal models. SDT has at its core the idea that observers only receive noisy representations of stimuli (Palmer et al., 1993; Green & Swets, 1966). The observer combines these noisy representations into a single variable, and then applies a threshold to this variable. If the variable exceeds a threshold, the observer reports the target is present, if not, absent.

SDT models predict graded changes in performance. Consider what happens as distractors become less similar to the target. The chance that noise will cause them to be confused with the target decreases, decreasing the false-alarm rate and therefore potentially increasing performance (Rosenholtz, 2001). In this respect, SDT models may make similar predictions to the claims of Duncan and Humphreys (1989). However, SDT models may also make contrasting predictions. Specifically, when the distractors and target are very similar, increasing distractor variability will spread the distractors out, away from the target. This will decrease the chance that they will be confused with the target, decreasing false alarm rate, and potentially increasing performance (Rosenholtz, 2001; see Fig. 1). Note that this mechanism may only be important on trials when the target is absent: When the target is present, the stimulus which looks most like the target is likely to be the target itself. Therefore, spreading distractors out may rarely make a difference to the stimulus which appears closest to the target.

**Figure 1:**
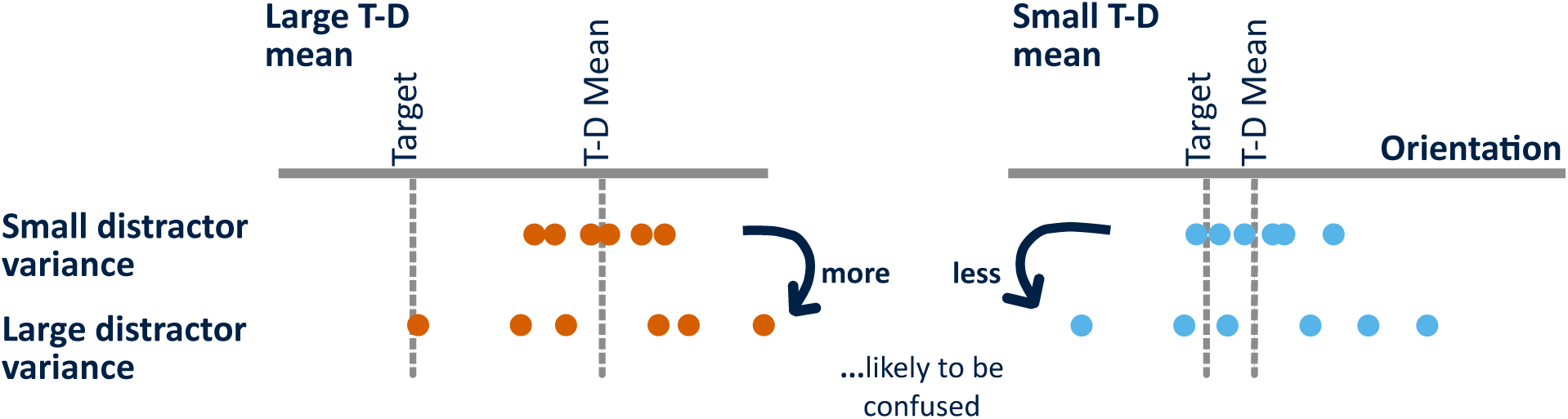
Interaction between target-to-distractor mean difference (T-D mean), and distractor variance on the probability of a confusing distractor. When T-D mean is large, then increasing distractor variance makes a distractor which closely resembles the target more likely. On the other hand, when T-D mean is small, increasing distractor variance actually makes it less likely that there is a distractor which is highly similar to the target.

Using process models, such as SDT models, we can generate predictions for the relationship between any stimulus statistic and any behavioural statistic. This is because these models predict behaviour on a trial-by-trial basis using the full stimulus, rather than a summary of the stimulus. Hence, we can always ask what the model predicts for stimuli low or high on any particular statistic. By contrast, if our theory is that distractor mean predicts accuracy in a certain manner, it remains completely unclear how other distractor statistics might predict accuracy, or how mean might be related to another behavioural statistic such as hit rate. For example, low accuracy could be caused by completely random responding, or by always picking the same response. It should be noted that Duncan and Humphreys (1989) developed a detailed account of how parts of a visual scene are grouped and compete for entry into visual short-term memory. They used this account to explain the effects of distractor statistics that they described. In this paper, we do not attempt to convert the entirety of their underlying theoretical account into a process model, but instead focus on the effects of distractor statistics.

Signal detection theory encompasses a range of approaches to visual search, of which the Bayesian approach is one (Green & Swets, 1966; Rosenholtz, 2001; Palmer et al., 2000). In the Bayesian approach, we assume that the observer computes a very specific single variable from the noisy stimuli representations, namely the posterior ratio. This is the ratio of the probability that the target is present and the probability that the target is absent, given the observer’s measurements (Palmer et al., 2000; Peterson, Birdsall & Fox, 1954; Ma et al., 2011). We assume that the observer has learned the statistical structure of the task, and computes the posterior ratio using this knowledge. This assumption results in a highly constrained model which has been shown to fit behaviour well in a range of visual search tasks (Ma et al., 2011; Mazyar, van den Berg, Seilheimer & Ma, 2013; Mazyar, van den Berg & Ma, 2012). However, the consistency of Bayesian observer models with the qualitative predictions of Duncan and Humphreys (1989) has not been examined in detail.

The present work has several goals. (a) We aim to describe the effect of target-distractor difference, and distractor variability across the full stimulus space, comparing the results with the claims of Duncan and Humphreys (1989), and with SDT ideas (see Fig. 1). (b) We examine whether a Bayesian optimal-observer model accounts for patterns in the data, and whether the Bayesian model is consistent with the claims of Duncan and Humphreys (1989). (c) We examine whether simpler heuristic models can also account for the observed patterns. Rosenholtz (2001) conducted closely related work, aiming to test the idea that, under SDT, increasing distractor variability might actually improve performance when the target-distractor difference is low. Rosenholtz (2001) only used specific stimulus values, ones ideal for the question of that paper. As a consequence, Rosenholtz (2001) did not broadly explore the stimulus space. Mazyar et al. (2013) conducted a very similar experiment to ours, but focused on the effects of number of stimuli on precision, rather than the effects of distractor statistics on performance.

A subtle but sufficiently important point to warrant discussion at the outset, is that we study one kind of distractor statistics. We explore the effects of statistics of sampled distractors. This contrasts with examining the effects of population statistics – the statistics of the distributions from which distractors are drawn. In the “Theory of Visual Selection” described by Duncan and Humphreys (1989, p.444), both sample and population distractor statistics have a role. We focused on sample distractor statistics because experimental study of these effects is more feasible. To study the effects of population distractor statistics, participants would need to be trained on many different distractor distributions. We trained participants on two distributions here but, to anticipate our results, it is unclear whether participants even learn the difference between these two distributions.

## 2 Experimental Methods

### Participants

14 participants were recruited consistent with a pre-determined recruitment schedule (see appendix A for age, gender and handedness information). One participant was excluded from all analysis below as they were unable to complete all sessions. The study procedure was approved by the Institutional Review Board of New York University and followed the Declaration of Helsinki. All participants gave informed consent.

### Apparatus

Stimuli were presented on an LCD monitor at 60 Hz refresh rate, with 1920 × 1080 resolution, and a viewable screen size of 51 × 29 cm. Stimuli were presented using Psychtoolbox in MATLAB on Windows (Brainard, 1997; Pelli, 1997; Kleiner, Brainard & Pelli, 2007). A chin rest ensured participants viewed the stimuli at approximately 60 cm from the screen. Eye tracking data was collected for future exploratory analysis, however, to date this data has not been analysed in any form.

### Stimuli

Each stimulus was a Gabor patch with a standard deviation of the Gaussian window of 0.25 degrees of visual angle (dva), and with 1.79 cycles per dva. Stimuli were presented on a grey background (half the maximum of the RGB range). At the centre of the Gaussian window, the peaks and troughs of the sinusoid were represented as greys at the maximum and minimum of the RGB range.

6 stimuli locations were determined at the start of the experiment. These locations were equally spaced around the circumference of an imagined circle. Therefore, in the plane of the screen, Gabors were 60 degrees apart from each other (the first patch was at 90 degrees from vertical). Each location was at 4.99 dva from the imaginary line connecting the participant to the centre of the screen. In trials where there were fewer than 6 stimuli to display, a subset of the locations were randomly selected.

### Trial procedure (Fig. 2)

Each trial began with a fixation cross presented at the centre of the screen for 500 ms. Participants were presented with 2,3, 4, or 6, Gabor patches for 100ms, and asked to report the presence or absence of a target. Participants had unlimited time to respond “target present” with the ‘j’ key, or “target absent” with the ‘f’ key.

**Figure 2:**
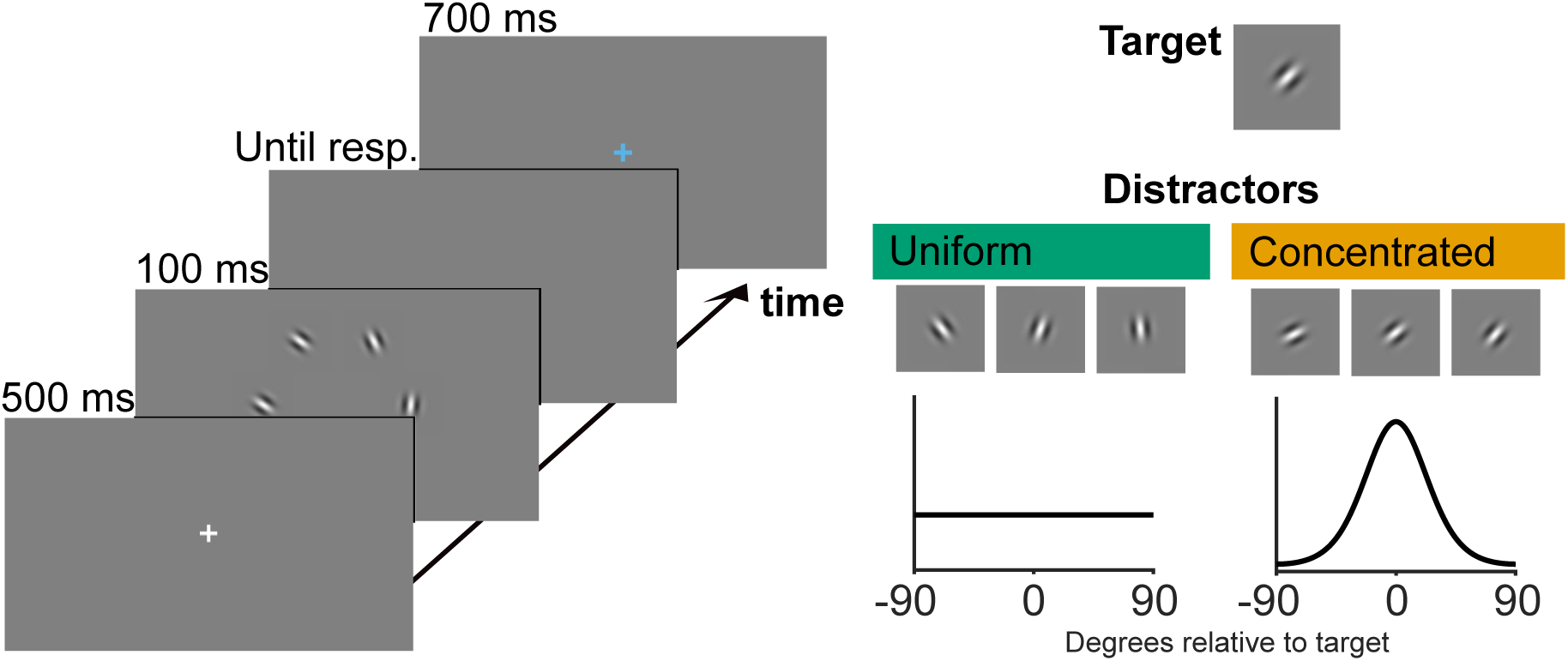
Participants performed a visual search task in which they had to report the presence or absence of a target (a Gabor oriented at 45 degrees clockwise from vertical) in a briefly presented display. The display contained between 2 and 6 stimuli. There were two distractor environments, one in which all distractor orientations were equally likely, and one in which distractor orientations were more likely to be close to the target orientation. “Screenshots” are for illustration, and not to scale.

The target was a Gabor patch oriented at 45 degrees clockwise from vertical. Targets were present on 50% of trials. All other Gabors in the display, named distractors, were drawn from a probability distribution which depended on the distractor environment (described below). It was possible for a distractor to have almost the same orientation as the target orientation.

Following a response, participants received feedback in the form of an orange or light blue fixation cross. This colour coded fixation cross was presented for 700 ms seconds. Following a trial there was a delay of at least 100 ms for setting up the eye tracker. The next trial would not begin until all keys had been released.

### Structure of the experiment

There were two distractor environments, and these determined the probability distributions from which distractors were drawn (Fig. 2). In the *uniform distractor environment*, distractors were drawn from a uniform distribution, and hence any orientation was equally likely. In the *concentrated distractor environment*, distractors were drawn from a von Mises distribution centred on the target orientation and with concentration parameter 1.5. The von Mises distribution is similar to the normal distribution, but is the appropriate choice for circular variables (i.e. orientation). The randomly drawn angles were divided by 2 to determine the orientation of the Gabor. This was done because a Gabor pointing up is identical to a Gabor pointing down. We can avoid this ambiguity by only using the angles between −90 and 90 degrees. Each block either contained trials from the uniform distractor environment, or trials from the concentrated distractor environment.

The experiment took place over 4 separate one-hour sessions. In each session, there were 8 test blocks of 64 trials. Uniform and concentrated distractor environment blocks were ordered as AABBBBAA. Whether “A” corresponded to the uniform or concentrated distractor environment was determined randomly at the beginning of each session.

### Training

At the beginning of each session, the participant was presented with an image of the target. Beside this image was a series of example distractors from the uniform distractor environment, followed by a series of example distractors from the concentrated distractor environment. At the beginning of the first session, the participant also completed 4 training blocks. At the beginning of subsequent sessions, they completed 2 training blocks. Each training block contained 40 trials. Uniform and concentrated blocks alternated, with the first block being selected randomly at the beginning of each session (matching the first test block). During test blocks, every time the distractor environment switched, the participant was presented with a refresher, in the form of another series of example distractors drawn from the upcoming distribution.

### Analysis

Throughout the paper, unless otherwise stated, we analyse the effect of distractor sample statistics. That is, on each trial we calculate distractor statistics using the distractors which were actually presented. Throughout the paper, target-to-distractor mean difference (T-D mean) will be used to refer to the absolute difference between the circular mean of the distractors and the target orientation. Distractor variance will refer to the circular variance. Definitions of circular mean and variance are provided by Berens (2009). Minimum target-distractor difference (min T-D difference) refers to the absolute difference between the target orientation, and the distractor orientation closest to the target orientation (Mazyar et al., 2012). For all circular statistics we used the CircStat toolbox (Berens, 2009).

Prior to computation of circular statistics, we double all orientations. The reason for this is that stimuli at −90 degrees are identical to those at 90. So that this is accounted for, we double orientations meaning that the new mapped orientations take up a full 360 degrees. In all plots, we map orientations (including T-D mean and min T-D difference) back to physical orientation.

In order to test the reliability of observed trends, we performed logistic regressions using distractor statistics as predictors, and either hits, false alarms (FA), or accuracy as outcome. (We included a constant as a predictor in each logistic regression.) We compared the fitted regression slopes to zero across participants. Prior to running the regression, we z-scored the predictors. Centring the variables allows interpretation of a main effect in the presence of an interaction, as the effect of the predictor at the mean value of all other predictors (Afshartous & Preston, 2011). We provide adjusted p-value significance thresholds, using the Bonferroni correction, to account for the number of regression slopes compared to zero in each individual regression analysis. As a measure of effect size we computed the one-sample variant of Cohen’s d (Cohen, 1988),

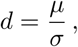

where *µ* is the mean of the beta values being compared to zero, and *σ* is the estimated population standard deviation.

For this analysis (and not for computational modelling below), data from trials with only two Gabor stimuli was excluded. The reason for this is that when there are only two stimuli and one of them is a target, there is only one distractor, and the idea of distractor variability does not make sense. Throughout the paper, unless labelled, plots reflect data from trials with 3, 4, and 6 stimuli.

### Plots

In order to visualise the effect of distractor statistics, we binned these variables. Specifically, we used quantile binning, separately for the data from each participant. We took this approach as distractor statistic distributions can be highly non-uniform (Fig. 3), and quantile binning ensures a reasonable number of data points in each bin. In order to determine where on the x-axis to plot a bin, we computed for each participant the average value in each bin, and then averaged these across participants. The location of a bin on the y-axis was determined by the mean value of the outcome variable across participants. Unless stated, error bars represent ±1 standard error of the mean.

**Figure 3:**
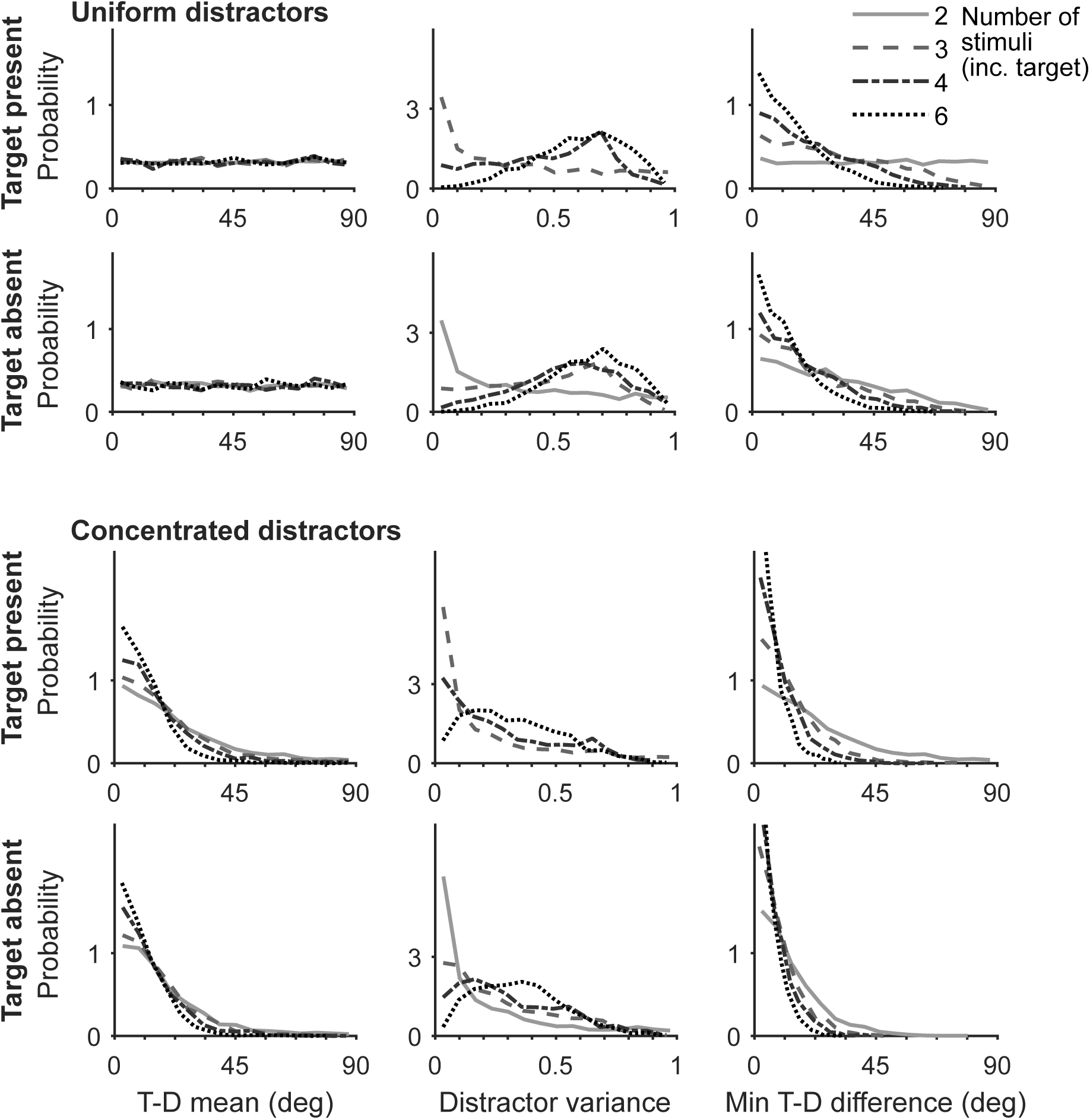
The distributions of distractor statistics, separately for the cases of 2, 3, 4, and 6 stimuli in the display (including the target). The area under the curves in all 12 plots is the same. Note that these distributions are determined by stimuli properties, and are completely independent of participant behaviour. Here and throughout the paper, target-to-distractor mean difference (T-D mean) refers to the absolute difference between the circular mean of the distractors and the target orientation, and distractor variance refers to the circular variance of the distractors. Minimum target-distractor difference (min T-D difference) refers to the absolute difference between the target orientation, and the distractor closest to this orientation. Data from all participants combined is shown. We can see that distractor statistic distributions are highly non-uniform. This is the motivation for, in all other plots than this one, quantile binning distractor statistics.

### Data and code availability

Anonymised data, together with all experiment and analysis code written for the study, will be made available upon publication at doi:10.17605/OSF.IO/NERZK.

## 3 Experimental Results

The first aim of the study was to explore the effect of distractor statistics using stimuli which could be parametrically varied. This allows fine-grained variation in distractor statistics and comprehensive exploration of the stimulus space. We explored these effects as participants reported the presence or absence of a Gabor oriented at 45 degrees clockwise from vertical. We turn to computational modelling of the identified patterns in later sections.

In an initial set of analyses we focused on testing the pattern of effects suggested by Duncan and Humphreys (1989). For each participant, we conducted regressions with target-to-distractor mean difference (T-D mean), distractor variance, and their interaction as predictors, and either accuracy, hit rate or false alarm (FA) rate as outcome. The resulting regression coefficients reflect the strength of the relationship between the predictor and the outcome. We compared these coefficients to zero (Fig. 4, and Table 1). T-D mean, distractor variance and their interaction significantly predicted accuracy. At the average value of distractor variance, increasing T-D mean increased accuracy, while at the average T-D mean, increasing distractor variance decreased accuracy. The two interacted such that at large T-D mean, the relationship between distractor variance and accuracy was more negative. This finding contradicts the idea of Duncan and Humphreys (1989) that increasing the heterogeneity of distractors (increasing distractor variability) would have relatively little effect on performance when target and distractors are very different from each other (high T-D mean). Whilst these effects are significant, they are difficult to observe directly from the plot (particularly the effect of distractor variance; Fig. 4, A). The small T-D mean series appears to exhibit a “U” shape, with the lowest accuracy values at a distractor variance of about 0.35. If real, this effect represents a systematic deviation from a logistic relationship, an assumption of using logistic regression. Therefore the result of this analysis should be interpreted with caution.

**Table 1:** The effect of T-D mean, distractor variance, and their interaction, in the absence of additional predictors. When no other predictors are included in the model all three variables have a significant effect on accuracy, hit rate, and FA rate, with the exception of the effect of the interaction on hit rate.

| Outcome | Predictor | t-Value | Effect size (d) | p-Value |
| --- | --- | --- | --- | --- |
| Accuracy | T-D mean | 5.5 | 1.5 | $1.3 \times 10^{-4}$ |
| | Distractor variance | -3.1 | -0.86 | $9.4 \times 10^{-3}$ |
| | Mean-var interaction | -3.2 | -0.89 | $7.5 \times 10^{-3}$ |
| Hit rate | T-D mean | -5.5 | -1.5 | $1.3 \times 10^{-4}$ |
| | Distractor variance | 3.5 | 0.97 | $4.5 \times 10^{-3}$ |
|  | Mean-var interaction | 2.2 | 0.61 | 0.049 |
| FA rate | T-D mean | -8.0 | -2.2 | $4.0 \times 10^{-6}$ |
| | Distractor variance | 4.2 | 1.2 | $1.2 \times 10^{-3}$ |
| | Mean-var interaction | 6.2 | 1.7 | $4.5 \times 10^{-5}$ |

Bonferroni corrected p-value criterion 0.017.

**Figure 4:**
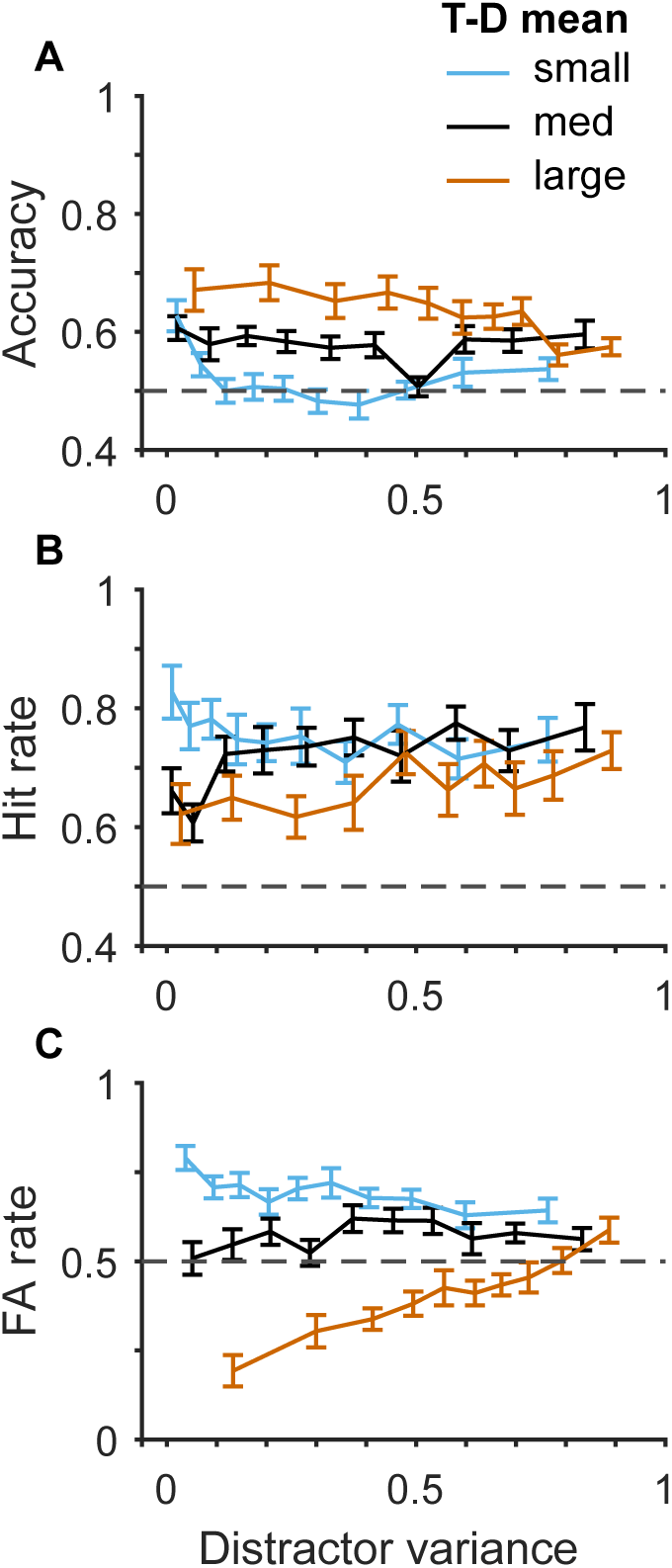
The effect of T-D mean and distractor variance on behaviour. There was a particularly clear interaction effect on FA rate, consistent with a signal detection theory account. For the plot T-D mean was divided into 3 bins, participant by participant. The average edges between bins were at 12 and 36 degrees. Data from trials with 3, 4, and 6 stimuli were used for plotting.

By contrast, T-D mean, distractor variance, and their interaction had a particularly clear effect on FA rate (Fig. 4, C). At the average value of distractor variance, increasing T-D mean decreased FA rate. At the average T-D mean, as distractor variance increased FA also increased. There was also an interaction such that distractor variance had the most positive effect for large T-D mean. This pattern is completely consistent with the pattern we would expect from a SDT perspective (recall Fig. 1). Specifically, for large T-D mean, increasing variance increases the probability of a distractor similar to the target. Whilst with a small T-D mean, increasing variance actually makes a confusing distractor less likely, as distractor orientations are spread out away from the target orientation.

A similar pattern of effects was observed on hit rate, although the pattern is harder to observe in this case (Table 1, Fig. 4, B). Note that the considerations above, regarding T-D mean and distractor variance interacting to affect the probability of a confusing distractor, do not directly apply in the case of hit rate. The hit rate is calculated using trials on which the target was in fact present, hence, the most similar stimulus to the target (from the perspective of the observer) is likely to be the target itself. This may explain why the effects of the distractors are diluted.

Having considered the effect of distractor mean and variance in isolation from other variables, we wanted to explore whether the effects identified could be due to variability shared with additional variables. For each participant, we used T-D mean, distractor variance, their interaction, the minimum target-distractor difference (min T-D difference), distractor environment, and number of stimuli, as predictors in logistic regressions to predict accuracy, hit rate or FA rate. As before, we compared the resulting coefficients to zero across participants. Only the min T-D difference and the number of stimuli significantly predicted accuracy, and only the min T-D difference predicted hit rate (Table 2). A possible explanation for the difference between these findings, and the regressions without additional variables included, is that it is the min T-D difference which is the causally relevant variable. The effects of T-D mean and distractor variance may appear because T-D mean and distractor variance are correlated with the min T-D difference. This suggests that while Duncan and Humphreys (1989) may have identified distractor statistics which are related to performance, they make not be the cause of changes in performance.

**Table 2:**
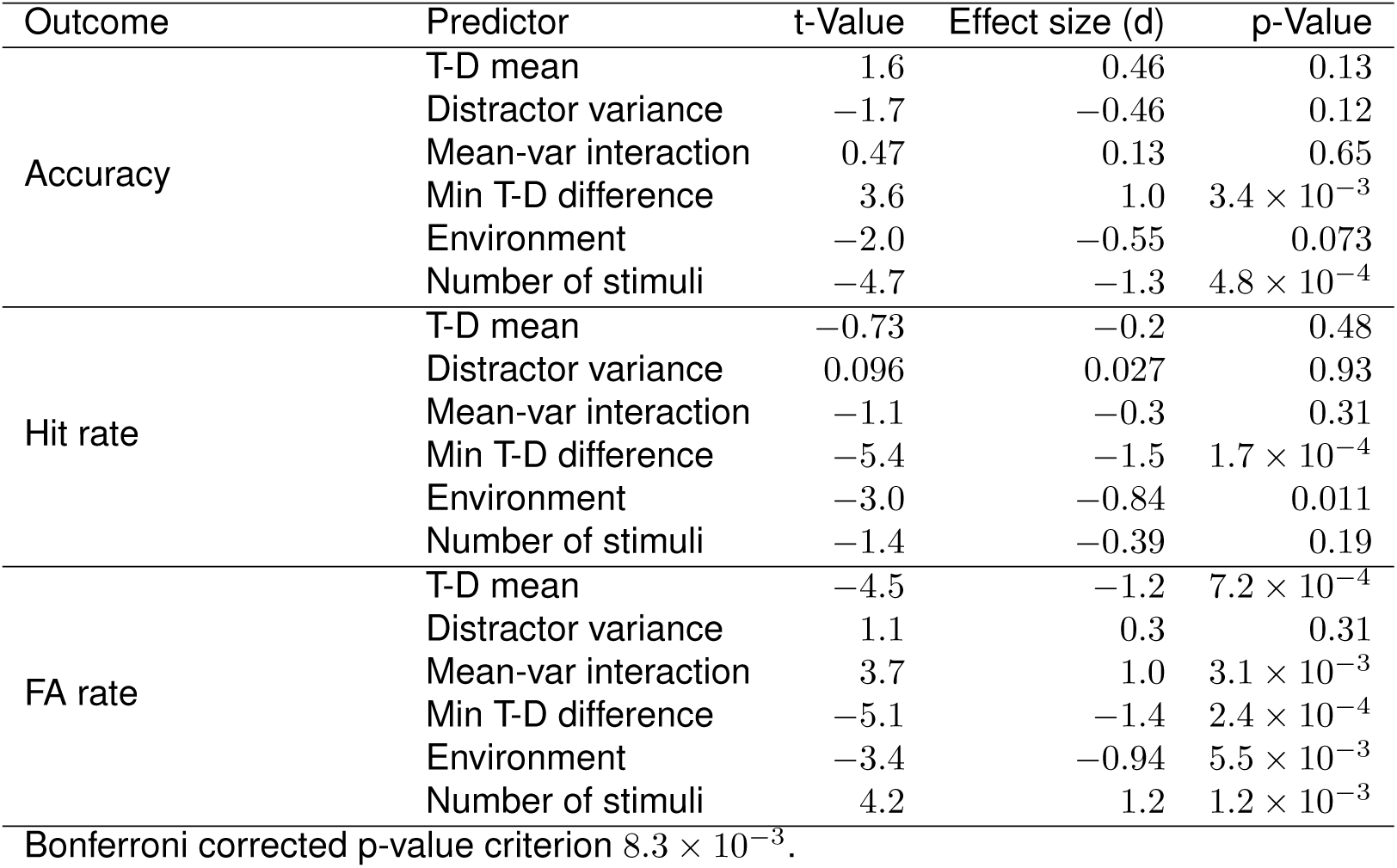
The effect of distractor and experiment variables on accuracy, hit rate, and FA rate. Surprisingly, T-D mean, distractor variance and their interaction do not have a significant effect on accuracy. Instead, the effect of min T-D difference is significant.

This finding that it is min T-D difference, not T-D mean or distractor variance, that matters for performance, may seem to conflict with previous work. Rosenholtz (2001) found that performance in visual search suffers when distractors are made more variable, even when this is done in a way that doesn’t move the distractors closer to the target. This study featured a blocked design, so participants may have successfully learned and used knowledge of the population from which distractors were drawn. It may be that variability in the population from which distractors are drawn is harmful to performance, even if variance of distractors in a particular sample is not.

A different pattern emerged when looking at the regression onto FA rate (Table 2). T-D mean and the interaction between T-D mean and distractor variance predicted FA rate in the same direction as they had without the inclusion of additional variables, although distractor variance was no longer a significant predictor. The min T-D difference had a large effect on FA rate, with FA rate decreasing as the min T-D difference increased. This finding provides evidence that T-D mean and distractor variance have some relevance to behaviour, over and above their relationship with min T-D difference. Note that this finding does not rule out the possibility that the most similar stimulus from the perspective of the observer, is the only important variable in determining their response. This is because the min T-D difference, according to the participant, will not necessarily be the true min T-D difference, due to perceptual noise. Therefore, other stimuli, not just the most similar, may affect behaviour even if the observer only uses the stimulus which appears most similar to them.

For the dedicated reader we provide univariate analyses of the effects of distractor statistics in appendix B.

To summarise, consistent with the account of Duncan and Humphreys (1989), there was evidence for an effect of T-D mean, and distractor variance. We detected an interaction effect between T-D mean and distractor variance, however, the effects of the interaction did not match the effects predicted by Duncan and Humphreys (1989). Plots suggested that the relationship between these variables and accuracy was not simple, and may not be well described by the regression model used. One of the advantages of building a process model of visual search is that complex patterns which evade adequate treatment with conventional statistics, may be accounted for in a parsimonious way. A similar pattern of distractor statistic effects was found on hit rate. The pattern of effects on FA rate was particularly clear and entirely consistent with SDT considerations. Analyses which also included the effects of additional variables suggested that, at least in the case of accuracy, min T-D difference is the causally relevant variable, not T-D mean or distractor variance.

## 4 Modelling Methods

We next explored whether a computational model could provide a parsimonious explanation of the effects identified. Of particular interest is whether a computational model can capture both patterns in the data which match those identified by Duncan and Humphreys (1989), and those which differ. We focus on a highly constrained Bayesian model initially, and compare this model to other models below. There are three steps to building an Bayesian observer model (Mazyar et al., 2012; Ma et al., 2011; Mazyar et al., 2013). First, we must specify how experiment stimuli are generated, and make assumptions about how information from the environment is encoded in the brain of the observer. Together this information forms the generative model, the model of the series of events that leads to the observer’s measurements. Second, we must specify the rule the observer uses to make decisions on the basis of these measurements. For an optimal observer, we must derive the optimal decision rule. Third, we must derive model predictions. This is an additional step because we do not have access to the observer’s measurements on a trial-to-trial basis. Instead, using our assumptions of how stimulus information is encoded, and knowledge of the stimuli presented, we can predict which measurements are more and less likely, and in turn, which responses are more or less likely.

### 4.1 Generative model

The first step is to specify how measurements are generated. The target is present on half of trials. Denote the presence of the target *C* = 1 and absence *C* = 0, then we have,

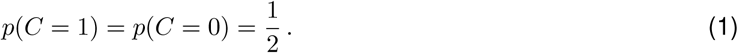

There are between 2 and 6 possible target locations, because there are between 2 and 6 stimuli in a display. If the target is in location *i* we write *T*_*i*_ = 1, and if it is absent in this location *T*_*i*_ = 0. **T** indicates a vector containing *T*_*i*_ for every location. If the target is present at location *i*, then this stimulus is at 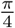 radians clockwise from vertical. We express this using the Dirac delta function,

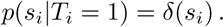

Here *s*_*i*_ represents the orientation of the Gabor at the *i*th location in radians. However, it represents the orientation in a very specific way. It represents twice the difference between the Gabor orientation and the target orientation. Therefore, to convert from *s*_*i*_, to orientation clockwise from vertical, in radians, *a*_*i*_, use,

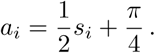

There are two reasons for using *s*_*i*_ rather than *a*_*i*_. First, it is convenient to have the target at *s*_*i*_ = 0. Second, as discussed in the methods, we only use Gabors between 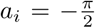 and 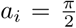 radians, to avoid the ambiguity caused by Gabors not having a direction. Using *s*_*i*_ simplifies mathematics as we don’t have to repeatedly write 2*a*_*i*_. As mentioned previously, in all plots, we map orientations (including T-D mean and min T-D difference) back to physical orientation. That is, we map all orientations back to the space of *a*_*i*_, not the transformed *s*_*i*_.

If the *i*th location contains not a target but a distractor, then the stimulus orientation is drawn from a von Mises distribution with mean *µ* = 0, and concentration parameter *κ*_*s*_,

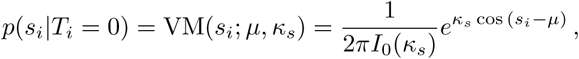

where VM indicates the von Mises distribution, and the second equality provides the definition of this function. *I*_0_ are modified Bessel functions of the first kind and order zero. A von Mises distribution is very similar to a normal distribution, but is the appropriate distribution for a circular variable (i.e. orientation). A von Mises distribution with *κ*_*s*_ = 0 is the same as a uniform distribution. Therefore, we can model both distractor environments, uniform and concentrated, with this equation.

Finally we assume that the observer only receives noisy measurements of the stimulus orientation. We formalise this by assuming that measurements are drawn from a von Mises distribution centred on the true stimulus orientation, but with concentration parameter *κ*,

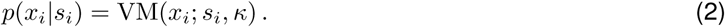

In words, we are assuming that measurements of stimulus orientations are variable, but unbiased, so that if you took the average of lots of measurements, you would almost recover the true stimulus orientation.

We can represent this model of how measurements are generated graphically (Fig. 5). In the figure and throughout, **s** and **x** in bold font represent vectors of *s*_*i*_ and *x*_*i*_ for all *i*.

**Figure 5:**
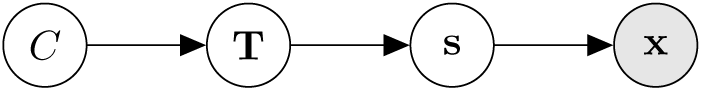
Representation of the generative model. The observer has to use their measurements, x, to infer whether or not the target is present *C*

### 4.2 Optimal decision rule

Having specified the generative model, we can now use Bayes’ rule to determine the optimal way to make decisions. Here we only state our premises and conclusion, but the full derivation is provided in appendix C.

Turning to our premises, we do not assume that the observer equally values hits and avoiding false alarms. Instead we include in our models a parameter, *p*_present_, which captures any bias towards reporting “target present”. We use the fact that there is at most one target, and assume that measurement noise at different locations is independent.

As shown in appendix C, from these assumptions we can derive the following rule for optimal behaviour. The observer should report “target present” when

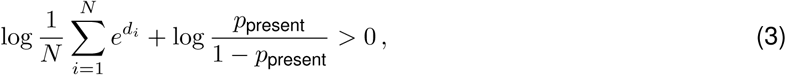

where

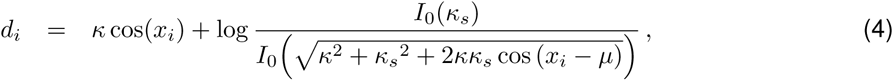

and *N* is the number of stimuli in the display.

### 4.3 Making predictions

We need to use our knowledge of how the observer makes decisions on the basis of **x**, to make predictions for how the observer will respond given **s**. If we denote the observer’s response *Ĉ*, then we want to find the probability *p*(*Ĉ*|**s**). By assumption, a stimulus, *s*_*i*_, generates measurements according to (2). Hence, for a particular set of stimuli, we can simulate measurements of these stimuli, and determine which responses these measurements lead to. By repeating this process many times we can build an estimate of the probability, according to the model, that a particular set of stimuli will lead to a “target present”, or “target absent” response. For each trial, we simulated 1000 sets of measurements and the associated decisions.

### 4.4 Lapses

We allow the possibility that some trials are the result of contaminant processes, such as getting distracted. On these “lapse” trials, the participant makes a random response. If we denote the probability of response, *Ĉ*, according to the Bayesian observer model without lapses, *p*_no lapse_(*Ĉ*|**s**), and the lapse rate *λ*, then the probability of a response is given by

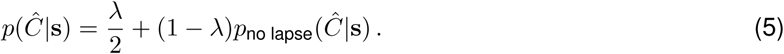

### 4.5 Model fitting

Separately for each participant, we fitted lapse rate (*λ*), bias parameter (*p*_present_), and the concentration parameter of measurement noise (*κ*) as free parameters. A greater value of *κ* means that the observer’s measurement of the stimulus is corrupted by less variability. We allow the possibility that measurement noise varies with the number of stimuli in the display, and fit *κ* as four free parameters (one for each possible number of stimuli in the display; Mazyar et al., 2013).

For any valid set of parameter values *θ*, we can calculate the likelihood. The likelihood is equal to the probability of the observed behaviour, given the parameters and the stimuli shown. Assuming that responses in different trials are independent of each other, we can write the likelihood as a product of the probability of responses on each trial,

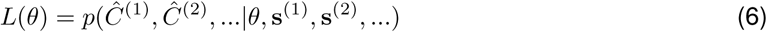

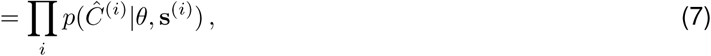

where the product is taken over all the trials for a participant, *Ĉ*^(*i*)^ is the participant’s response on the *i*th trial, and **s**^(*i*)^ denotes the stimuli on the *i*th trial. We used Bayesian adaptive direct search (BADS) to search for the parameters which maximised the log-likelihood (Acerbi and Ma, 2017). BADS is a well tested optimisation algorithm which alternates between a poll stage in which nearby parameter values are evaluated, and a search stage in which a Gaussian process model is fitted and used to determine promising parameter values to evaluate.

The model was fit separately for each participant. For each participant we ran BADS 40 times. For each run, 150 parameter value sets were randomly selected and the likelihood evaluated at each. The set with the highest likelihood was used as the start point for the run. The bounds on the parameters during the search, and the way in which initial parameter values were drawn, is described in appendix E. Running the fitting procedure many times reduces the chance of getting stuck in local maxima, and permits heuristic assessment of any problems local maxima may be causing (see supplementary methods of Acerbi, Dokka, Angelaki & Ma, 2018). We found that fits to the same log-likelihood function often ended at different values of “maximum” log-likelihood, suggesting that we may have only found local maxima, rather than finding the global maximum. For a discussion of these issues, and our attempts to resolve them by reducing noise in the likelihood function, see appendix F.

### 4.6 Alternative models

Given the strong effect of the minimum target-distractor difference on behaviour, a heuristic which focuses on the measured orientation most similar to the target orientation might perform well. We compared the Bayesian observer to an observer who uses a very simple decision rule. If the absolute difference between the measured distractor orientation closest to the target orientation, and the target orientation, is below some threshold *ρ*, they report “target present”, otherwise they report “target absent”. Models of this kind have been used extensively in visual search research (Ma, Shen, Dziugaite & van den Berg, 2015). Note that, because the observer applies a criterion to a noisy variable to determine their response, this heuristic observer model is also a SDT model (Palmer et al., 2000; Palmer et al., 1993). We make predictions for behaviour in the same way that we did for the optimal observer model, and fit the model in the same way.

We fitted two variants of this model. In model 3 (see Table 3) the threshold used by the observer varies with different numbers of stimuli in the display, and varies in different distractor environments. There are 4 possible numbers of stimuli in the display (2, 3, 4 and 6), and 2 distractor environments, giving a total of 8 thresholds which were fitted as free parameters. In model 4 we allowed the threshold to vary with number of stimuli in the display but assumed that it was fixed across distractor environments, as if participants ignored the difference between the environments when making their decisions.

**Table 3:** Parameters of all models considered in the paper. The main model is a Bayesian optimal observer (model 1). We also considered an observer which applied a heuristic, and made their decision entirely on the basis of the measured orientation closest to the target orientation (models 3 and 4).

| Model number | 1 | 2 | 3 | 4 |
| --- | --- | --- | --- | --- |
| Inference | Bayes | Bayes | Heuristic | Heuristic |
| Distractor environment use | True | False | True | False |
| **Parameters** |  |  |  |  |
| Sensory noise ($\kappa$) | 4 | 4 | 4 | 4 |
| Decision thresholds ($\rho$) | n/a | n/a | 8 | 4 |
| Bias ($p_{\text{present}}$) | 1 | 1 | n/a | n/a |
| Stimulus variability ($\kappa_o$) | 0 | 1 | n/a | n/a |
| Lapse rate ($\lambda$) | 1 | 1 | 1 | 1 |
| **Total** | **6** | **7** | **13** | **9** |

We also included a variant on the Bayesian observer model in our model comparison. The Bayesian observer discussed in the previous section is model 1, but we also consider a model in which the observer is Bayesian, except they ignore the difference between the two distractor environments. Instead this observer assumes all stimuli, regardless of distractor environment, are distributed following a von Mises distribution with a concentration parameter which we fit, *κ*_*o*_. This is model 2. A list of all parameters and models is shown in Table 3.

We used the Akaike information criterion (AIC), and the Bayesian information criterion (BIC) to compare the performance of these models. The AIC and BIC take into account the likelihood of the fitted models, and the flexibility of each model in terms of the number of fitted parameters. A lower AIC and BIC indicates better fit. For each information criterion, we found the best fitting model across participants using the mean value of the information criterion. To determine whether the difference in fit between the best fitting model and the other models was meaningful we calculated, for each participant, the difference in information criterion between the overall best fitting model and each of the other models. By bootstrapping these differences 10000 times we computed 95% confidence intervals around the mean difference between the best fitting model and each of the other models. If the confidence interval on the mean difference does not include zero for all competitor models, then we can conclude that the best fitting model fits better than all other models.

## 5 Modelling Results

We explore whether a Bayesian observer model can explain the effects of distractor statistics by fitting such a model, before simulating data using the parameter values fitted for each participant. The simulated data represents the model’s predictions for how participants would respond to stimuli. By plotting both the real data and model simulated data on the same plot, we can visually inspect whether the model successfully accounts for the trends in human behaviour. For plots we simulated 24000 trials per participant. In plots we use error bars for data, and shading for model predictions. Shading, like error bars, covers ±1 standard error of the mean.

We first looked at whether the model can successfully account for the individual effects of the distractor statistics. Looking at Fig. 6 we can see that the model predictions closely match the observed data, and that all qualitative patterns are recovered. Consistent with the data and with Duncan and Humphreys (1989), the model predicts increased performance with increasing target-to-distractor mean difference (T-D mean). In contrast to the account of Duncan and Humphreys (1989), but consistent with the data, the model does not predict a strong relationship between distractor variance and performance.

**Figure 6:**
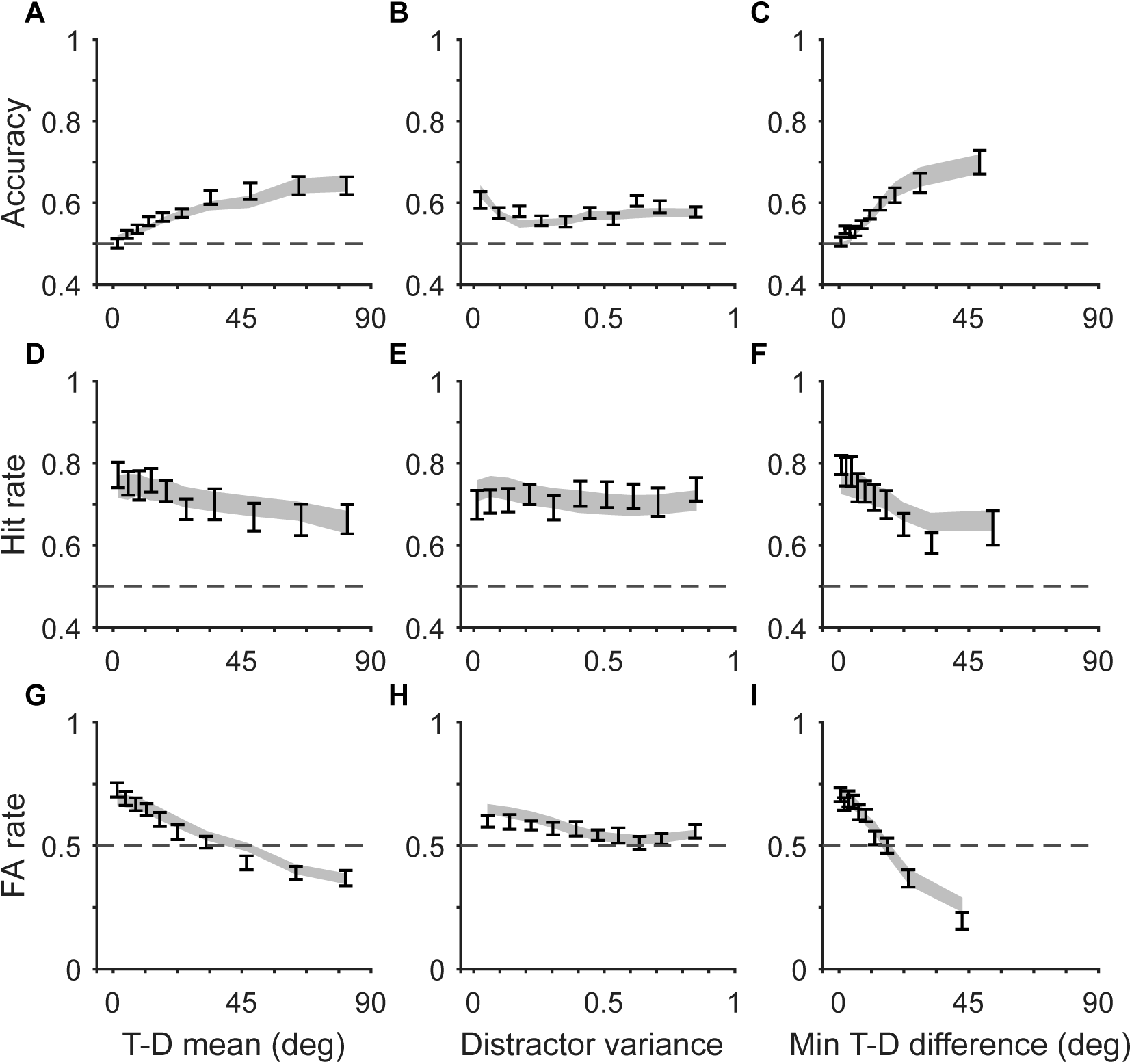
The effect of all summary statistics when considered individually (error bars), and model predictions for these effects (shading). The Bayesian observer model captures the observed effects well.

We noted in the experimental results that minimum target-distractor difference (min T-D difference) may be the causally relevant variable. The strength of the effect of min T-D difference is accurately captured by the Bayesian model (Fig. 6, C). It is interesting to ask why the Bayesian model would predict such a strong effect of just one distractor, when the Bayesian observer combines all distractor measurements in their decision rule (see equations 3 and 4). It turns out that, under specific conditions, the Bayesian observer closely approximates an observer who makes their decisions only on the basis of the stimulus which appears closest to the target. Fig. 7 shows the decision threshold the Bayesian observer applies to their measurements. The x- and y-axis represent measurements of two stimuli. If the measurements fall within the marked area, the observer reports “target present”. The decision thresholds are shown for a range of distractor environments, including the uniform and concentrated environments used in our study (concentration parameters *κ*_*s*_ = 0 and *κ*_*s*_ = 1.5). We can see that for distractor environments with more variable distractors (smaller values of the concentration parameter), the decision thresholds are such that, if any one of the two measurements is close to the target orientation the observer responds “target present”, regardless of the other measurement.

**Figure 7:**
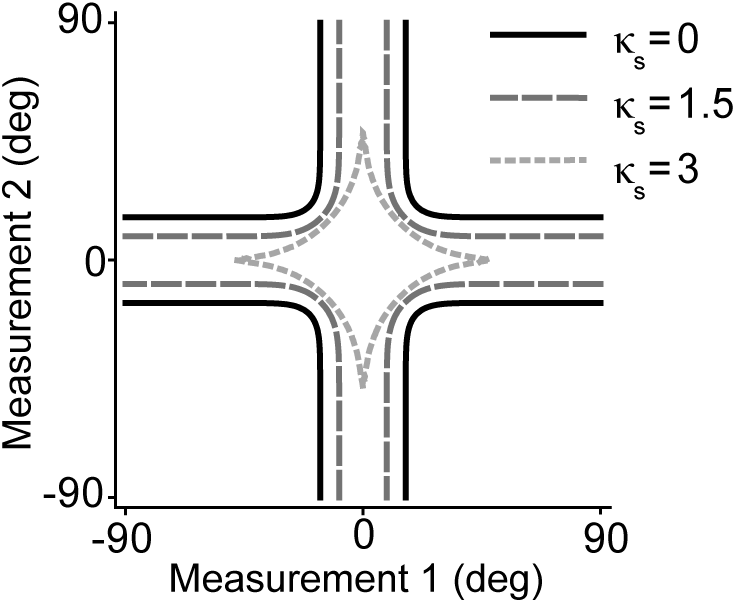
Decision thresholds used by the Bayesian observer for the case of two stimuli. The axes represent measurements of the stimuli made by the observer, relative to the target orientation. If the measurements fall within the marked area the observer reports “target present”. The thresholds were calculated using *κ* = 8, and under a range of values for *κ*_*s*_, including those used in the experiment (uniform environment, *κ*_*s*_ = 0; concentrated environment, *κ*_*s*_ = 1.5). For low *κ*_*s*_, the observer effectively only uses the measurement closest to the target to make their decision.

We next looked at whether the model could capture the observed interaction between T-D mean and distractor variance. The model captures the interaction between T-D mean and distractor variance on FA rate, including the decrease in FA rate with distractor variance at small T-D mean (Fig. 8, C). As discussed, this interaction on FA rate is predicted by signal detection theory accounts, because T-D mean and distractor variance have an interactive effect on the probability of a confusing distractor (Rosenholtz, 2001). The Bayesian observer, as a specific kind of SDT observer, inherits this effect.

**Figure 8:**
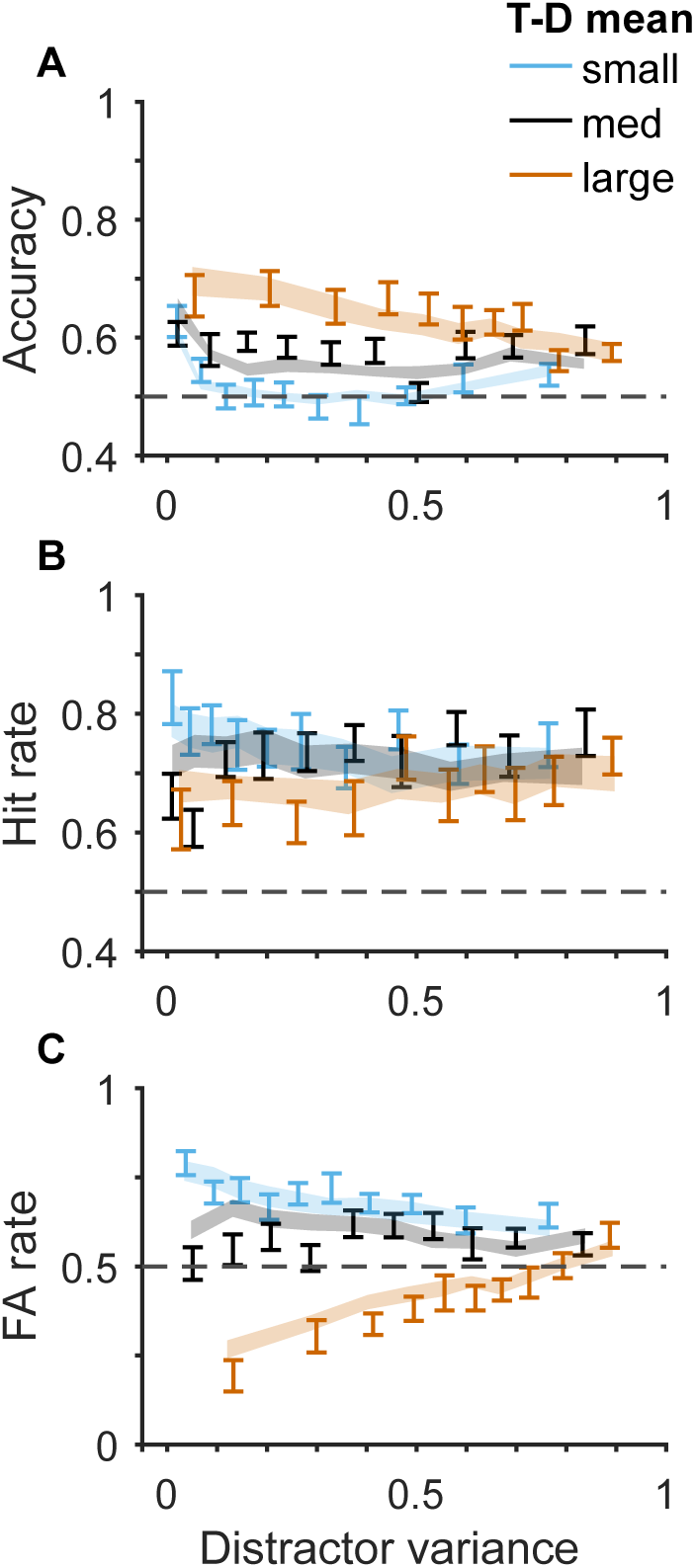
Interaction between T-D mean and distractor variance, and model predictions for these effects. The model captures the interaction of T-D mean and distractor variance on FA rate, along with the trends in accuracy.

The model also captures effects on accuracy and hit rate. In particular it accounts for the weak relationship between distractor variance, T-D mean and hit rate (Fig. 8, B). Looking at the model predictions for accuracy, we can see that the model largely captures the quantitative patterns (Fig. 8, A). Interestingly, the model captures the “U” shaped relationship between accuracy and variance for small T-D mean trials. This relationship would be overlooked using regressions or logistic regressions alone, but emerges out of an optimal observer model.

Much previous research has focused on the effect of number of stimuli in the display (e.g. Treisman and Gelade, 1980; Duncan and Humphreys, 1989). We examined whether the Bayesian model could account for the effects of number of stimuli on accuracy, hit rate and FA rate (Fig. 9). As in previous work (Mazyar et al., 2012; Mazyar et al., 2013), Bayesian observer model predictions were highly accurate. The model captured the reduction in accuracy with number of stimuli, the largely flat effect on hit rate, and the increase in false alarms. It captured the effect of number of stimuli in both distractor environments, and additionally, captured the difference between the environments.

**Figure 9:**
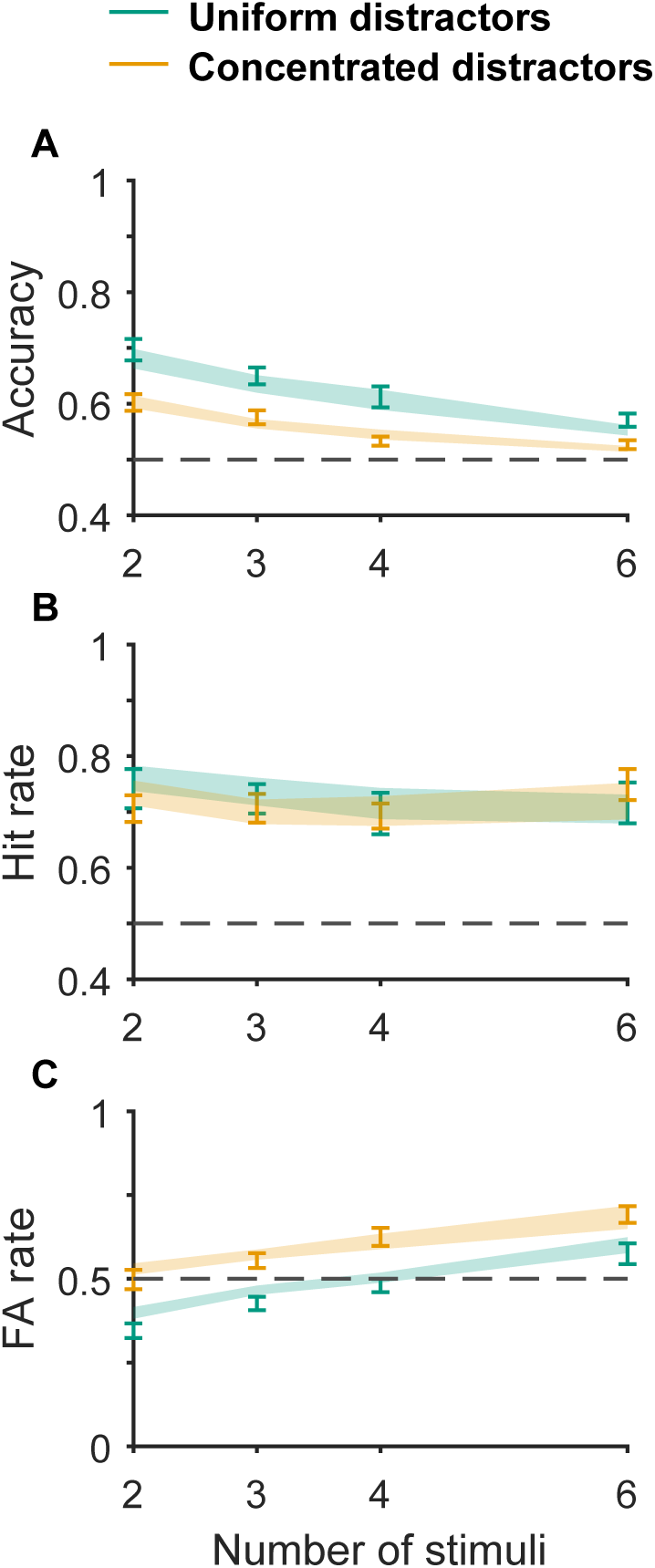
The effect of number of stimuli. The Bayesian observer model captured the reduction in accuracy with more stimuli, and the increase in false alarms.

Finally, we looked at whether the model could account for fine grained details, by looking at the effect of distractor statistics separately for different numbers of stimuli (uniform environment: Fig. 10; concentrated environment: Fig. 11). Mazyar et al. (2013) established that Bayesian observer models can account for the effects of min T-D difference in displays with different numbers of stimuli. We looked at these effects here, but also looked at the effect of T-D mean and distractor variance. The model largely captures the effect of distractor statistics for all stimuli numbers. We note that there appear to be some systematic deviations from the model predictions. For example, for two stimuli, and concentrated distractors, the model does not capture an apparent dip in hit rate at median values of min T-D difference (Fig. 11, C). If reliable, this is an intriguing phenomenon: When the most similar distractor is very different to the target, “target present” responses are more probable than when the most similar distractor is just somewhat different. A similar pattern has been observed before (Mazyar et al., 2012). This suggests that some part of the mechanism of visual search may not be captured by the Bayesian model.

**Figure 10:**
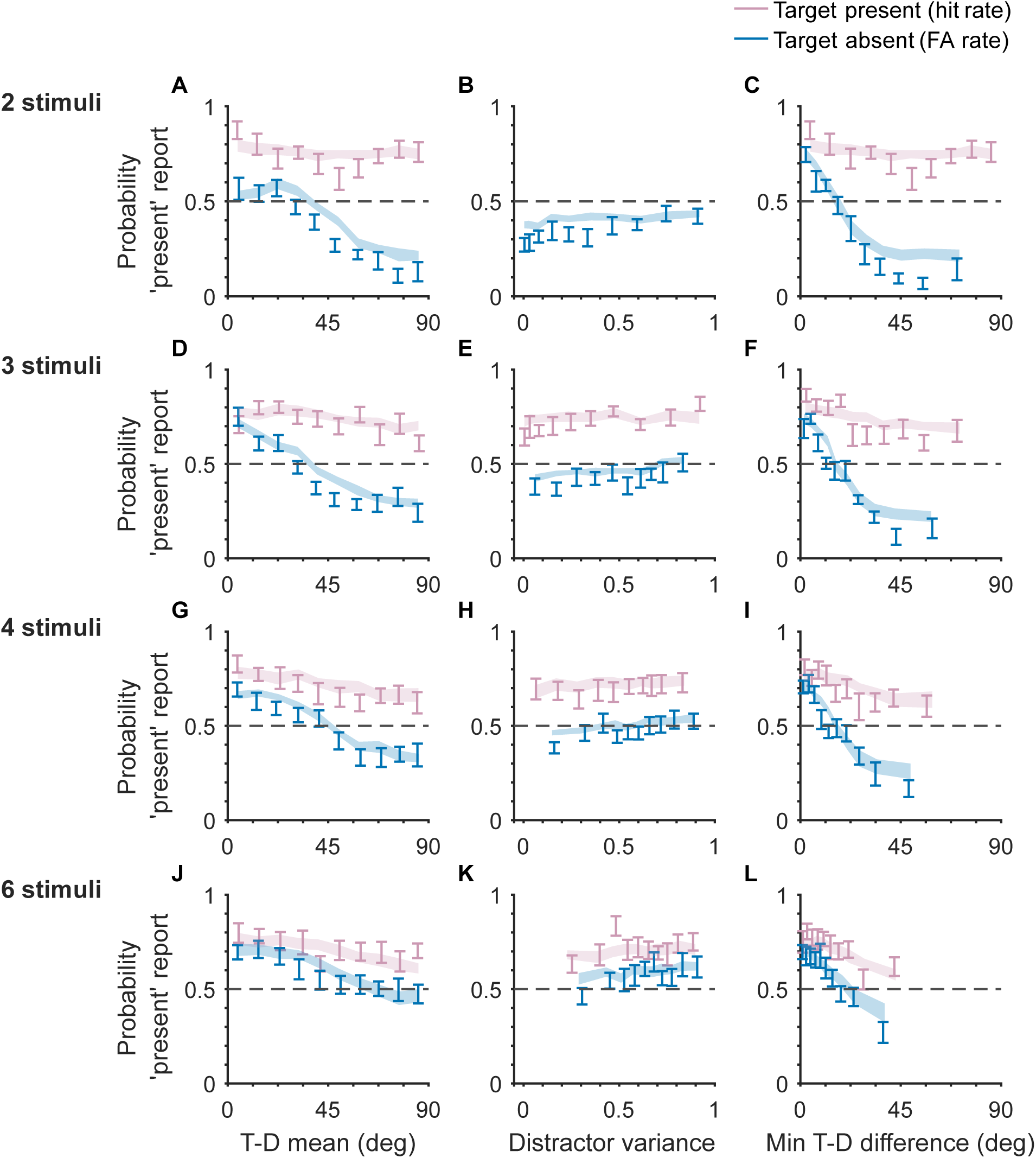
Effect of distractor statistics in the uniform environment, at different numbers of stimuli. The Bayesian observer model successfully accounts for most effects at all numbers of stimuli considered, although there appear to be some systematic deviations.

**Figure 11:**
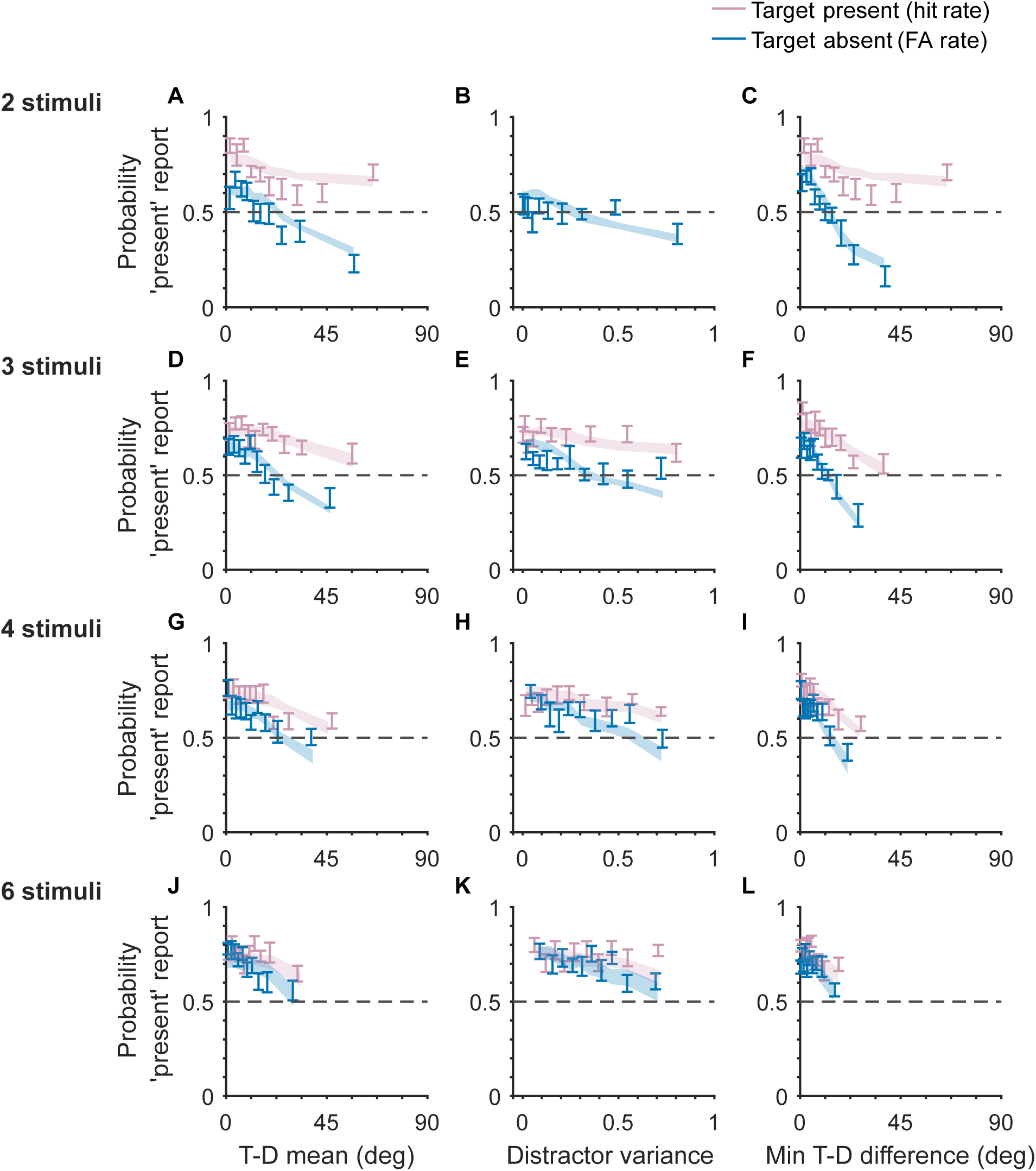
Effect of distractor statistics in the concentrated environment, at different numbers of stimuli. The Bayesian observer model successfully accounts for most effects observed.

Having seen that a Bayesian observer model captures trends in the data well, we wanted to explore whether other models could also explain the data as well or better. Of particular interest is the question of whether a model in which the observer uses a heuristic might explain the observed data better. We compared two heuristic observer models (3 and 4), to two Bayesian observer models (1 and 2). The heuristic observer applies a threshold on the distractor which appears most similar to the target (from their perspective), to determine their response.

The results of the model comparison are presented in Fig. 12. According to the AIC, a heuristic observer (model 3) fit best. Model 4 is similar to model 3, however, in model 3 the observer applies different decision thresholds depending on the distractor environment. Confidence intervals on the difference in fit between this and the other models did not include zero, suggesting model 3 fit reliably better according to the AIC. Fig. 12 also shows that, according to the AIC, a majority of participants were best fit by model 3. In contrast, according to the BIC, the Bayesian optimal observer (model 1) fit best. However, confidence intervals on the BIC differences suggested that the difference in fit between this model and the other models was not reliable. According to the BIC a majority of participants were best fit by model 1.

**Figure 12:**
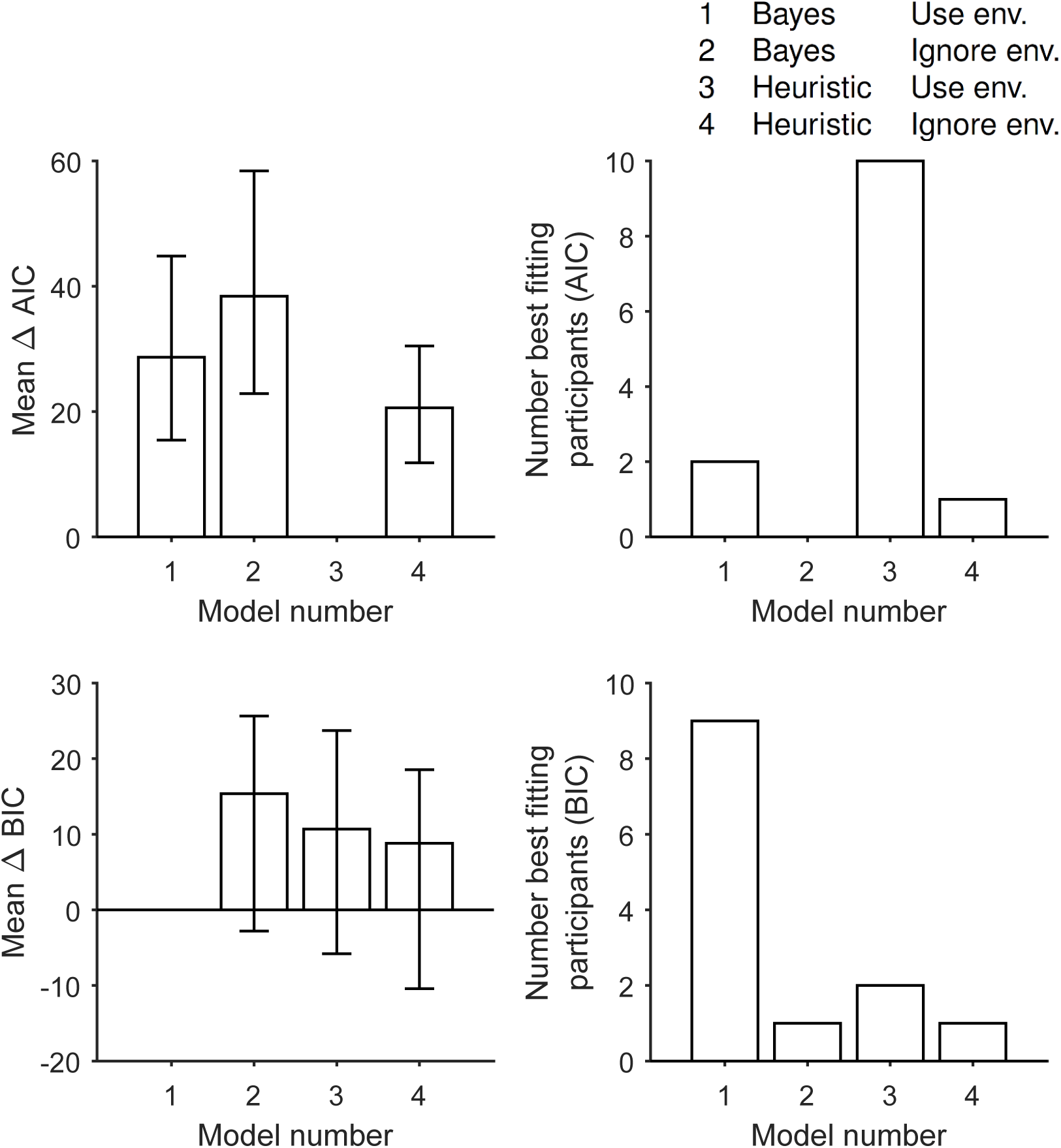
Mean AIC and BIC relative to the best fitting model, and the number of participants best fit by each model. See Table 3 for details of the models. Model comparison results were inconclusive because a consistent pattern of results was not found across AIC and BIC. Unlike in other plots, error bars here reflect 95% bootstrapped confidence intervals.

We do not want the conclusions of our research to depend on the fit metric used. Therefore, in this case we cannot draw conclusions about which model fit best. The differences between the AIC and BIC results stem from the fact that the BIC penalises extra model parameters more harshly. Model 3, best according to the AIC, has the most parameters out of all the models and so would be penalised heavily by the BIC (see Table 3). Using the AIC and BIC alone, we cannot say whether this penalisation is fair or not.

To explore these results further we performed model recovery analysis. We simulated data sets of the same size as the real data set using the participant-by-participant fitted parameter values. We then ran the above analysis on each of these simulated data sets. The only case in which the model used to simulate the data was not the best fitting model, according to the information criteria, was when data were simulated using model 3. On the AIC, model 3 fit best, as expected. However, according to the BIC, model 1 fit best. This suggests that in the present case, BIC may be unreasonably harsh on complex models. We note also, that the median fitted lapse rate for model 1 was 0.32, which seems unreasonably high (appendix D). In contrast, the median fitted lapse rate for model 3 was 0.15. Hence, there is very tentative evidence pointing to model 3 as the best model.

Bayesian observer models, and heuristic observer models of the kind considered here, have proved difficult to distinguish in previous work where the target takes a single value, and distractors are of equal reliability (Ma et al., 2015, sec. 2.3.2). We had hoped that the task used here, with two different distractor environments, could tease apart the models, but that proved not to be the case. The discussion above may help us understand why the Bayesian and heuristic observer models are difficult to distinguish: Under certain parameter values the Bayesian observer effectively only uses the measured orientation closest to the target orientation to make their decision, just as the heuristic observer does (Fig. 7).

We could also not decisively say whether observers used or ignored the difference between the distractor environments. (In model 2 and model 4, observers ignored the difference between the uniform and concentrated distractor environments; Table 3.) Both models which used, and those which ignored the difference between distractor environments, could predict different effects of distractor statistics in the two environments. For example, Fig. 13 shows data and model predictions for model 2, a Bayesian model in which the observer ignores the difference between the two distractor environments. In spite of this fact, the model predicts differences in the effect of distractor variance between the two environments. Such effects must be due to correlations between distractor environment, and other distractor statistics which do have an effect on behaviour.

**Figure 13:**
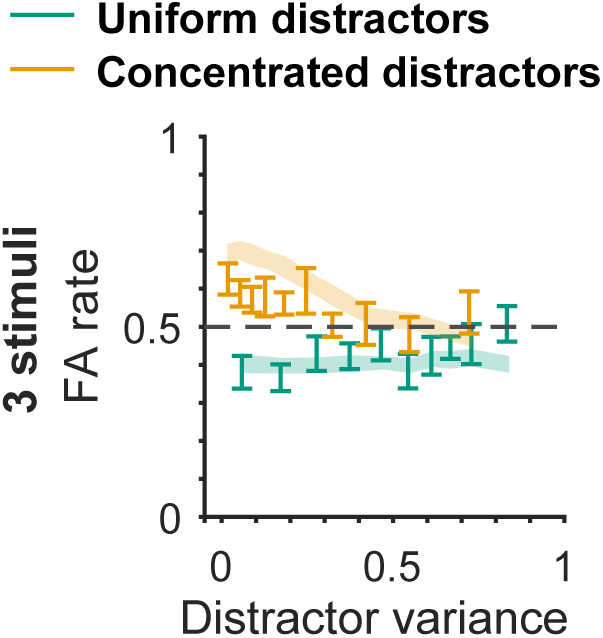
Data and model 2 predictions for the effect of distractor variance on FA rate. Model 2 assumes observers ignore the difference between the two distractor environments. Nevertheless, the model can predict differences between the two environments. This is likely because distractor environment correlates with other distractor statistics.

Parameter estimates from the fits are provided in appendix D.

## 6 General Discussion

In this study, we asked participants to perform a visual search task with heterogeneous distractors. Results regarding the distractor statistic effects identified by Duncan and Humphreys (1989) were mixed. We found some evidence for an effect of target-to-distractor mean difference (T-D mean), and distractor variance on accuracy. There was also evidence for an interaction between T-D mean and distractor variance on accuracy, but this interaction led to effects opposite to those predicted by Duncan and Humphreys (1989). We found that a statistic not explicitly considered by Duncan and Humphreys (1989), namely minimum target-distractor difference (min T-D difference), had a strong effect on behaviour, and that the effects of T-D mean and distractor variance may in fact be consequences of the effect of min T-D difference.

One potential reason for the discrepancy between our results and the account of Duncan and Humphreys (1989) is that we explored the difficulty of visual search through accuracy, whilst their primary variable of interest was response time (to be precise, the increase in response time as the number of stimuli increased). Specifically in the context of visual search, there is evidence that stimuli which generate low accuracy also generate slow responses (Eckstein, Thomas, Palmer & Shimozaki, 2000; Palmer, 1998; Geisler & Chou, 1995). Hence, difference in primary variable seems an unlikely explanation of the discrepancy between our results and the account of Duncan and Humphreys (1989). Nevertheless, it would be worthwhile to explore our data with a process model which makes predictions for response time as well as accuracy.

A second potentially important difference between our work and the work of Duncan and Humphreys (1989) involves the calculation of distractor statistics. In the present work, we explored the effects of statistics of sampled distractors. Duncan and Humphreys (1989, p.444) held that both sample and population distractor statistics (statistics of the population from which distractors are drawn) have a role. Studying the effect of population distractor statistics would involve training participants on a wide range of probability distributions. We attempted to train participants on two stimuli distributions (uniform and concentrated distractors). As discussed, we could not decisively say whether participants learned and utilised the difference between environments, suggesting training participants on a wide range of distributions would be challenging. As an alternative, future studies could explore the effects of population statistics using large numbers of participants and a between-participants design. It remains possible then, that the patterns identified by Duncan and Humphreys (1989) accurately describe the effects of population distractor statistics.

In the second half of this paper, modelling revealed that a Bayesian model with only 6 free parameters could account for a rich pattern of effects of distractor statistics. It captured the way T-D mean, distractor variance and min T-D difference, affected accuracy, false alarm (FA), and hit rate. It also accounted for the interaction between T-D mean and distractor variance, various effects of set size, and effects of distractor statistics at different set sizes. A model comparison of the Bayesian model with a variant, and with a heuristic observer model was inconclusive. This may be due in part to the similarity of the decision rule for the Bayesian and the heuristic observer: The Bayesian observer may effectively only use the stimulus which looks most similar to the target (Fig. 7), the policy of the heuristic observer.

Whilst we were unable to determine which model fit the data best, our findings suggest that SDT models (of which the Bayesian observer model and heuristic observer model are variants) can provide parsimonious explanations for a large set of distractor statistic phenomena. This work highlights some of the advantages of computational modelling. In particular, by building a process model of how stimuli are mapped to response, we were able to make predictions for a very wide range of effects. In fact, we could make predictions for how any distractor statistic affects any statistic summarising behaviour.

These findings complement other work showing that SDT models can provide parsimonious explanations of apparently complex phenomena in visual search. For example, SDT models can account for the apparent distinction between feature and conjunction search discussed in the introduction. This apparent distinction emerges from an SDT model that makes sensible assumptions about how multi-dimensional stimuli are encoded, but the model does not need to treat feature and conjunction searches as qualitatively different processes (Eckstein et al., 2000).

Our work has a number of limitations in scope, stemming from the fact that the experimental setup was chosen to ensure the experiment was well controlled and that modelling of behaviour was tractable. Perhaps of most importance is that stimuli were only presented for 100 ms, precluding the possibility of saccades. Preventing saccades is a common experimental choice (e.g. Eckstein et al., 2000; Palmer et al., 2000; Mazyar et al., 2013); the rationale is that it limits the complexity of the system under investigation. Specifically, we can ignore processes such as saccade selection and integration of previously gathered information, and instead focus on encoding and decision (see Table 4). Nevertheless, much visual search research has focused on tasks in which viewing time is unlimited. Importantly for the present discussion, many of the experiments in Duncan and Humphreys (1989) featured unlimited viewing time. Discrepancies between our results and the work of Duncan and Humphreys (1989) might, therefore, stem from the effects of processes studied by Duncan and Humphreys (1989), but not here. For instance, distractor variability might have a negative effect on the quality of saccades selected.

**Table 4:** Naturalistic visual search involves a large number of processes. In the present study, through the design of the experiment we focused on processes (b) and (c).

| Component processes in visual search | Example study |
| --- | --- |
| a Isolating objects from background | Wolfe, Alvarez, Rosenholtz, Kuzmova and Sherman (2011) |
| b Encoding of sensory information | Shen and Ma (2019) |
| c Decision mechanism | This paper |
| d Saccade selection | Najemnik and Geisler (2005) |
| e Integration of information from previous saccades | Horowitz and Wolfe (1998) |

Another decision which may limit the scope of the results is the use of relatively simple search displays. The stimuli used were easily distinguished from the surround, from each other, varied only along a single dimension, were not correlated with each other, and were limited to small numbers in each display (Palmer et al., 2000; Bhardwaj, van den Berg, Ma & Josić, 2016). Researchers have sometimes used many more than six stimuli (e.g. Treisman & Gelade, 1980; Rosenholtz, 2001), although using approximately six is certainly not an unusual choice (e.g. Duncan & Humphreys, 1989; Palmer et al., 2000). It is possible that the effects identified here would not generalise to tasks used in previous visual search work with many more stimuli. On the other hand, in many studies using a large number of stimuli, distractors have only taken one of a small set of values (e.g. Treisman & Gelade, 1980). In a important sense, the complexity of the stimuli used here is greater: the stimuli could take an infinite number of values, and all stimuli in the display took different values to each other. It would be premature then, to dismiss the conclusions of the present study on the grounds that the task used is simpler than tasks in previous research.

Even more important than whether the results can be compared to previous research is the question of whether the results generalise to naturalistic visual search. Palmer et al. (2000) highlighted many ways in which naturalistic visual search differs from conditions in lab studies. In real-world visual search, targets are unlikely to take specific values (e.g. you want to detect any car or motorcycle approaching, not just one specific car), vary along a signal dimension (e.g. cars vary in lots of ways), appear at a fixed location (e.g. a car could be anywhere along a road), involve small numbers of stimuli or be presented against a plain background (e.g. cars will be in a scene with signs, pedestrians, houses, and trees). By using simple, briefly presented stimuli, we have clearly not studied all the processes involves in naturalistic visual search (Table 4).

As scientists, our shared aim is to build a complete understanding of visual search, not just as it operates in the lab, but in naturalistic settings. Nevertheless, there are definite advantages to studying component processes separately. The choice of stimuli in the present study allowed us to explore the encoding and decision mechanisms of visual search in isolation. Thus, this choice vastly simplified the research problem, and increased the chances of producing intelligible results. We have seen in this paper just how successful our models of single stages (here the decision stage) can be. Moreover, there is much research on the other processes which make up visual search (see Table 4). The present paper contributed to this shared effort to understand naturalistic visual search by demonstrating that models of the decision mechanism provide an excellent account of the effects of distractor statistics.

## 7 Acknowledgements

We would like to thank Andra Mihali and Heiko Schütt for helpful discussions and advice over the course of the project. We are very grateful to Luigi Acerbi for extensive advice on dealing with model fitting difficulties. Wei Ji Ma was supported by grant R01 EY020958-09 from the National Institutes of Health. We would like to acknowledge the use of the University of Oxford Advanced Research Computing (ARC) facility in carrying out this work. http://dx.doi.org/10.5281/zenodo.22558

## A Participant demographics

**Table 5:** Aggregated gender, age, and handedness information for participants in the study.

|  |  |  |
| --- | --- | --- |
| Gender | Female | 10 |
|  | Male | 4 |
|  | Non-binary | 0 |
|  | Prefer not to say | 0 |
| Age | 18-25 | 10 |
|  | 26-35 | 3 |
|  | 36-45 | 1 |
|  | 46-55; 56-65; 66+ | 0 |
| Handedness | Right | 10 |
|  | Left | 3 |
|  | Neither | 1 |

## B Univariate analysis of distractor statistic effects

In addition to the analysis discussed in the main text, we also looked at the effect of distractor statistics when considered individually (i.e. ignoring variance shared with other distractor statistics and experimental variables). For each participant, we used target-to-distractor mean difference (T-D mean), distractor variance, or the minimum target-distractor difference (min T-D difference) in a logistic regression to predict accuracy, hit rate or FA rate. We compared the regression coefficients to zero across participants.

As expected, if the mean of the distractors was further from the target orientation, participants were less likely to report “target present” (Fig. 14, D and G; Table 6). Increasing T-D mean also increased accuracy of responses (Fig. 14, A; Table 6). Surprisingly, distractor variance was only related to FA rate. Increasing distractor variance predicted fewer false alarms (Fig. 14, H; Table 6). Like T-D mean, the min T-D difference strongly predicted accuracy and hit and FA rate. As the min T-D difference increased, the probability of a “target present” report decreased (Fig. 14, F, I; Table 6). At the same time, accuracy increased (Fig. 14, C; Table 6).

**Table 6:** The effect of distractor statistics on accuracy, hit rate and FA rate, when these effects are considered independently of the effects of other distractor statistics (including the other distractor statistics in this table), and experiment variables.

| Outcome | Predictor | t-Value | Effect size (d) | p-Value |
| --- | --- | --- | --- | --- |
| Accuracy | T-D mean | 6.1 | 1.7 | $5.5 \times 10^{-5}$ |
| Accuracy | Distractor variance | 0.22 | 0.061 | 0.83 |
| Accuracy | Min T-D difference | 5.0 | 1.4 | $3.3 \times 10^{-4}$ |
| Hit rate | T-D mean | -5.1 | -1.4 | $2.7 \times 10^{-4}$ |
| Hit rate | Distractor variance | 1.7 | 0.48 | 0.11 |
| Hit rate | Min T-D difference | -8.8 | -2.4 | $1.5 \times 10^{-6}$ |
| FA rate | T-D mean | -8.9 | -2.5 | $1.3 \times 10^{-6}$ |
| FA rate | Distractor variance | -3.0 | -0.84 | 0.011 |
| FA rate | Min T-D difference | -7.0 | -1.9 | $1.4 \times 10^{-5}$ |

**Figure 14:**
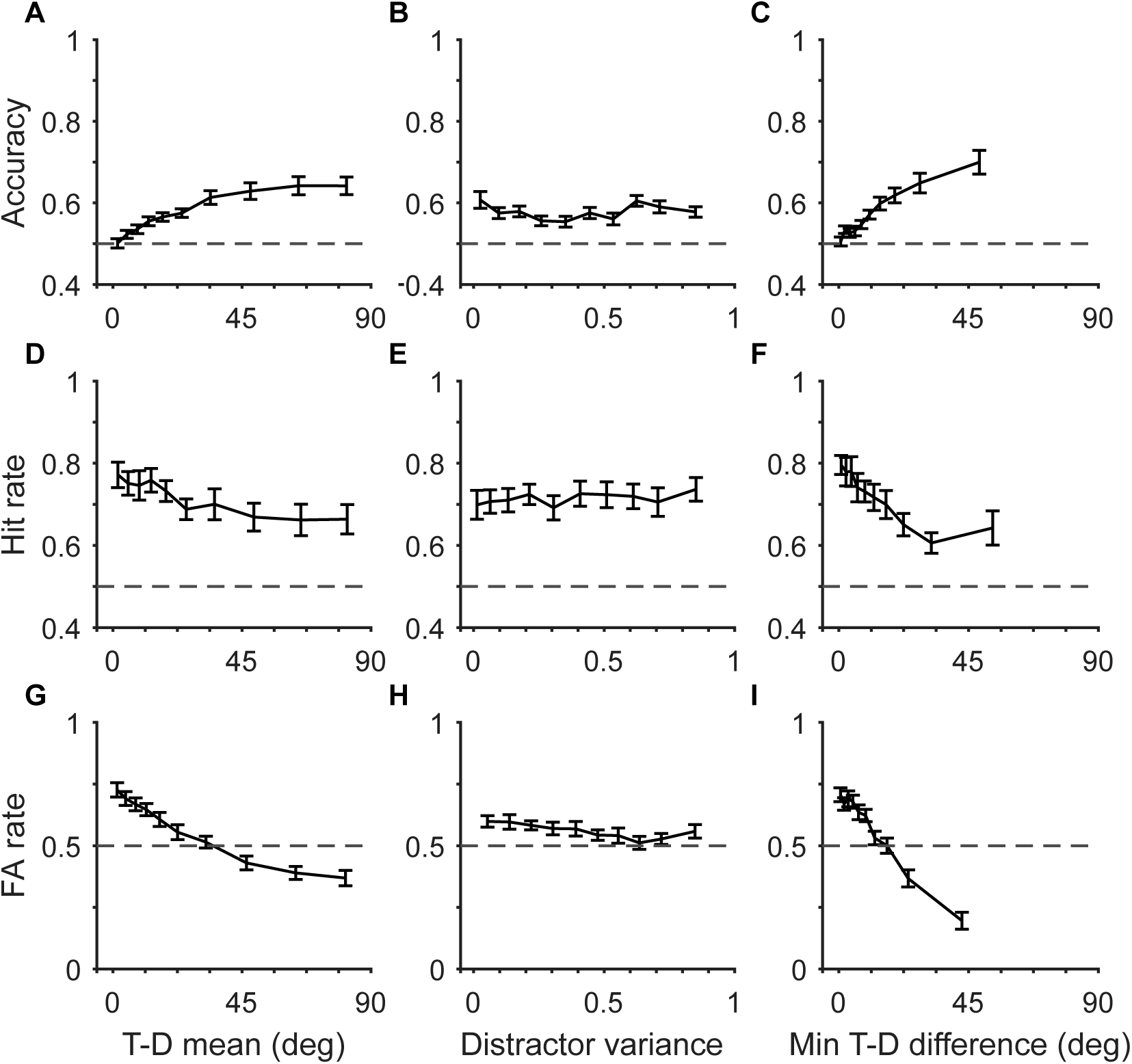
The individual effect of three distractor statistics on accuracy, hit rate and FA rate. As the T-D mean and min T-D difference increased, performance also increased and “target present” responses decreased. Surprisingly, distractor variance only had an effect on FA rate.

The effects of T-D mean and min T-D difference on “target present” responses suggest that participants were using a sensible strategy to perform the task: If the distractors were less like the target, participants were less likely to report “target present”. In addition, we observed that with increasing T-D mean and min T-D difference, performance improved. This finding suggests that similarity of target and distractors is an important determinant of performance. The lack of an effect of variance on performance may be because, as discussed in the introduction, increasing variance can make easily confused distractors either more or less likely, depending on the value of the T-D mean.

## C Derivation of the optimal decision rule

A Bayes-optimal observer uses the true generative model and evidence received in the form of measurements, to infer the probability of “target present” and “target absent”. An observer who equally values hits and avoiding false alarms, responds that the target is present when this is more likely than the target being absent. This is the same as using a criterion of 1 on the ratio of these probabilities, or 0 on the log of this ratio. We can write the condition for responding “target present” as,

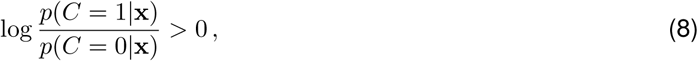

where *p*(*C* = 1|x) is the probability of the target being present after incorporating the information provided by the measurements x (the posterior probability).

We will not assume that the observer equally values hits and avoiding false alarms. Instead, much like in SDT, allow for the possibility that the observer values hits more than avoiding false alarms, or vice versa. Hence, in our models we will use the following decision rule,

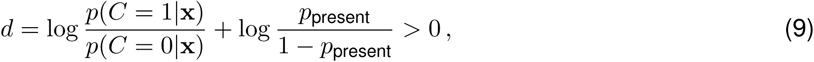

where *p*_present_ is the parameter which captures any bias towards reporting “target present”, and *d* denotes the sum of the posterior ratio and the bias term. Using Bayes’ rule and taking logarithms, we have

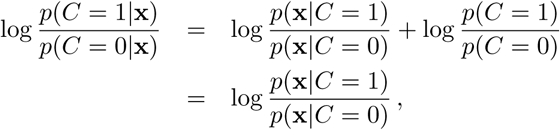

where the second line follows from equation (1). Hence, the optimal observer will report “target present” when

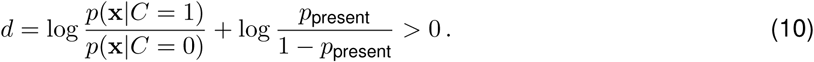

Assuming that there is at most one target, and that measurement noise at different locations is independent, it has been shown that the log-likelihood ratio is given by (Ma et al., 2011; Palmer et al., 2000)

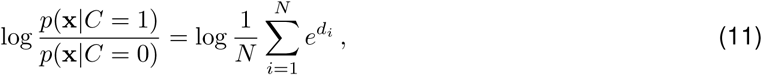

where *N* indicates the total number of Gabor patches in the display, and *d*_*i*_ indicates local log-likelihood ratio for location *i*. The local log-likelihood ratio is defined as

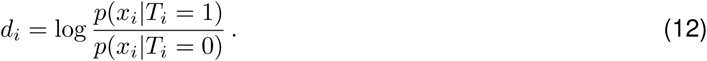

Marginalising over *s*_*i*_ and substituting in expressions from the generative model, we find

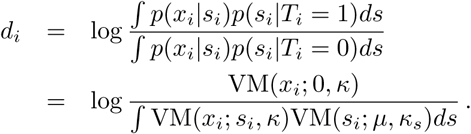

The denominator in this expression is the product of two von Mises distributions. Murray and Morgenstern (2010) state that the product of two von Mises is a new, scaled, von Mises. Any von Mises distribution, integrated over all angles, gives 1, because it is a probability distribution. Hence, when we integrate over all *s*_*i*_ we will only be left with the scaling. Using the formula from Murray and Morgenstern (2010) we have,

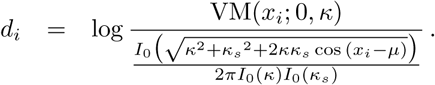

Substituting in the definition of a von Mises distribution and rearranging we find,

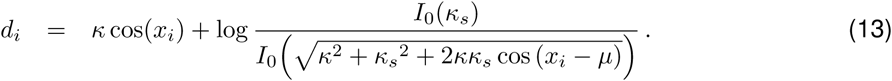

For the case of uniform distractors, *κ*_*s*_ = 0, and we have,

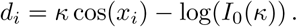

Substituting these expressions into equation (11) will give us, the log-likelihood ratio. In turn, using the log-likelihood ratio in (10) gives us the optimal observer’s decision rule. That is, it tells us, for any combination of measurements **x**, what the optimal observer would do.

## D Parameter estimates

Across participants we computed the median parameter estimate for each parameter, along with the 25th and 75th percentile.

**Table 7:**
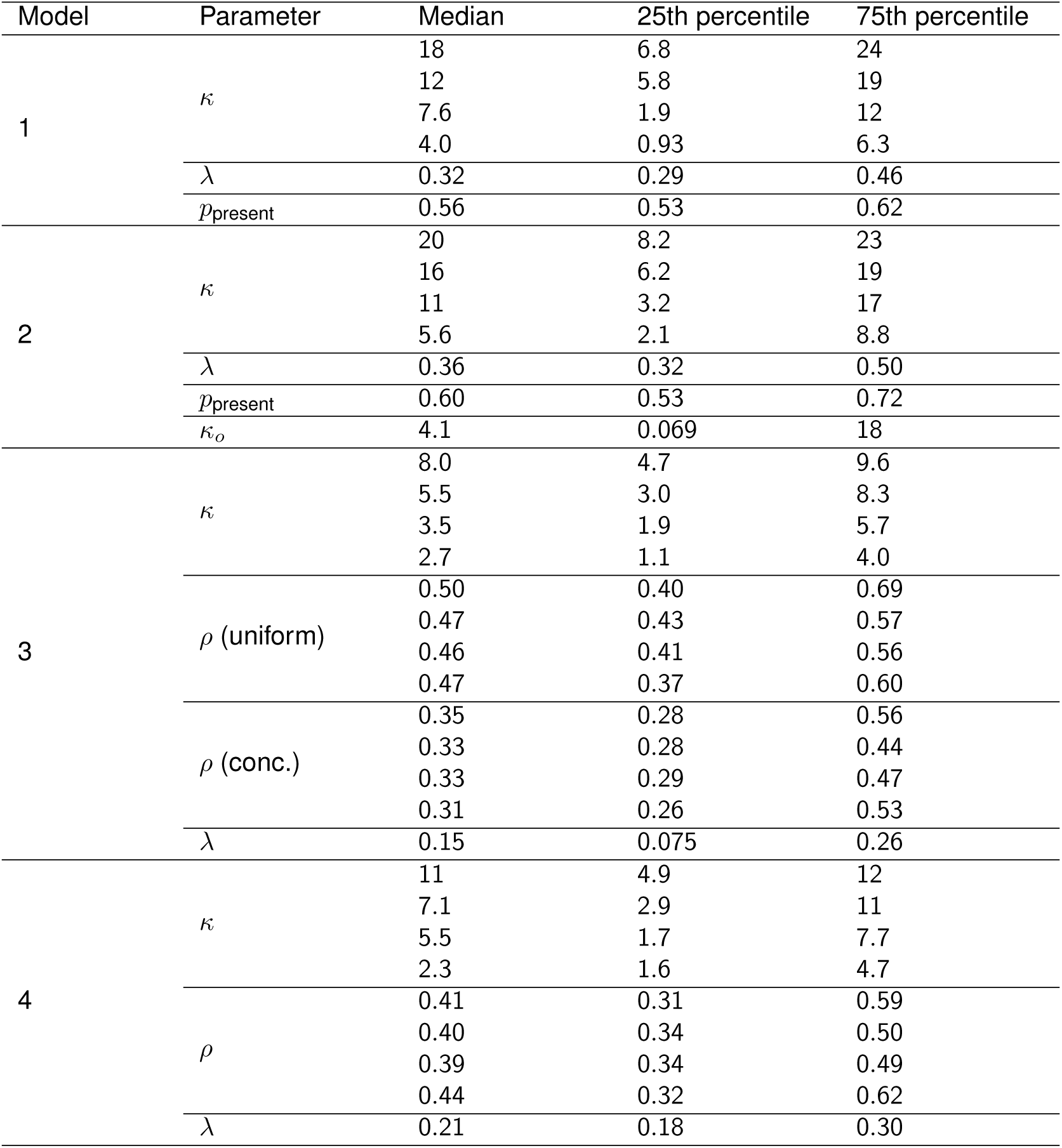
Parameter estimates. *ρ* is specified in radians.

## E Parameter bounds

During fitting with Bayesian adaptive direct search (BADS), we applied bounds to the values that the parameters could take, and specified plausible bounds within which we expected to find the parameter values. The bounds are specified below.

**Table 8:** Parameter bounds. *ρ* is specified in radians.

| Parameter | Lower bound | Upper bound | Plausible lower bound | Plausible upper bound |
| --- | --- | --- | --- | --- |
| $\kappa$ | 0.02 | 50 | 0.1 | 15 |
| $\lambda$ | 0 | 1 | 0.005 | 0.4 |
| $\rho$ | 0 | $\pi$ | $0.02 \pi$ | $0.75 \pi$ |
| $p_{\text{present}}$ | 0 | 1 | 0.2 | 0.8 |
| $\kappa_o$ | 0 | 100 | 0.2 | 25 |

Initial parameter values were drawn from uniform distributions on the interval between the plausible lower and upper bounds. For *κ* and *ρ*, sets of initial values were drawn until a set was drawn in which the parameters decreased monotonically with increasing number of stimuli on the screen.

## F Problems with local maxima

For each model and participant, we performed maximum-likelihood fitting 40 times. Fitting 40 times allowed us to estimate the probability that our best fits were reaching the true maximum-likelihood, as oppose to getting stuck in local maxima (see supplementary methods of Acerbi et al., 2018). To do this, we looked at how many fits ended up close to the best value of the likelihood found. If few fits get close to the maximum found, this suggests that the optimisation algorithm is struggling, and the global maximum may not have been found. For two models (2 and 3) many fits did not end up close to the best value of the likelihood (Fig. 15A), suggesting problems with fitting, and that an even greater value of the likelihood might exist but have been missed.

**Figure 15:**
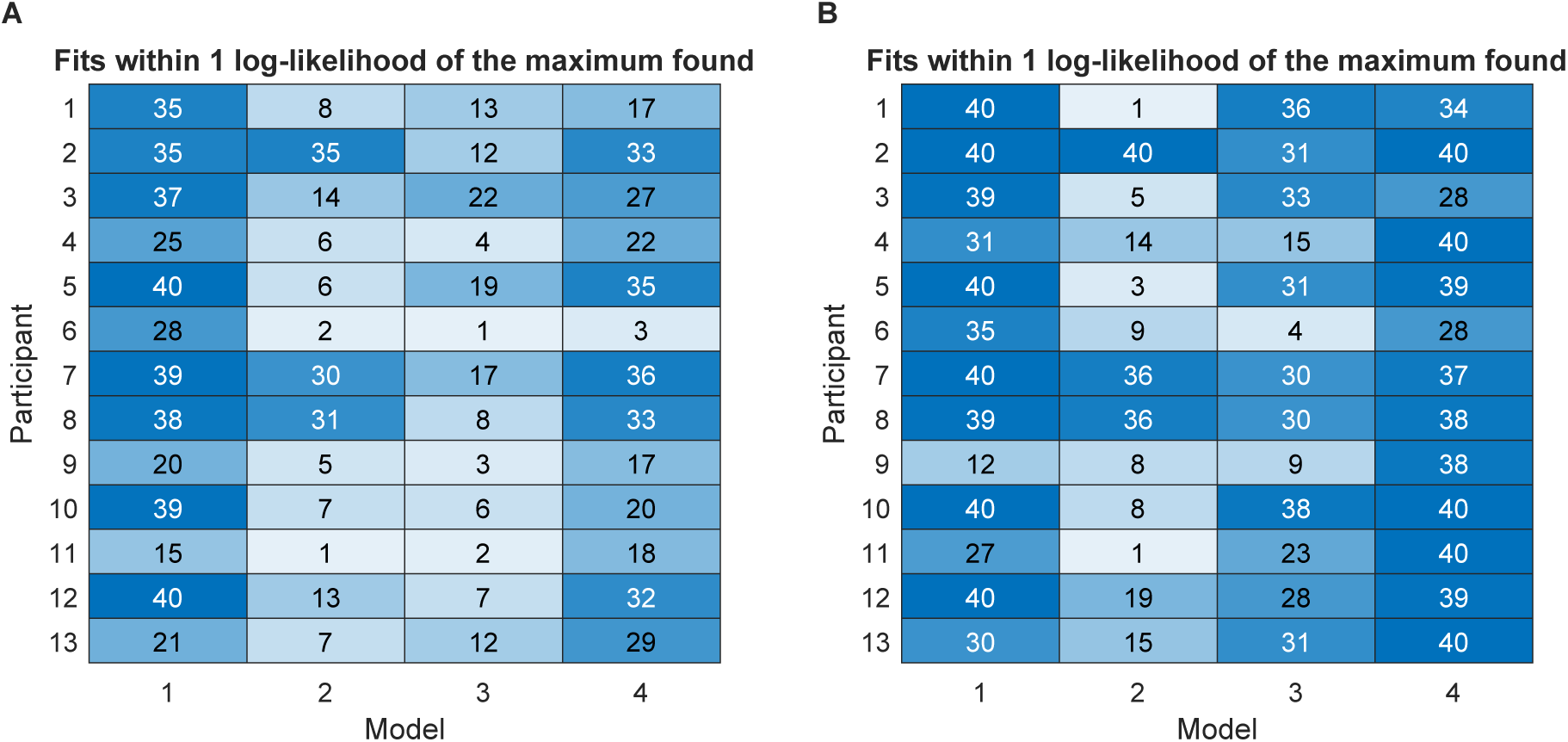
Number of fits out of 40 resulting in a log-likelihood within 1 point of the maximum log-likelihood found. Model numbers refer to Table 3. More saturated colors represent higher success rates. **A.** First run. **B.** Second run. The differences between the runs are described in the text.

To explore the possibility of issues with local maxima, we ran a further 40 fits for each model and participant, starting from the 40 points found in the first round of fitting. To reduce noise in the likelihood function, we simulated 10000 sets of measurements and associated decisions per trial, instead of 1000 as before. To make this approach computationally tractable, each likelihood evaluation we drew 5000 samples from each von Mises distribution (one for each value of *κ*), and re-sampled with replacement from these when von Mises samples were required. This improved the number of fits ending close to the best likelihood found for model 3, although there was still some evidence of potential issues with model 2 (Fig. 15B).

Since the results of the model comparison remained largely unchanged, we report the results for the first 40 fits in the main text.

